# A multi-resolution imaging and analysis pipeline for comparative circuit reconstruction in insects

**DOI:** 10.64898/2026.02.25.708065

**Authors:** Valentin Gillet, Marcel E. Sayre, Griffin S. Badalamente, Nicole L. Schieber, Kevin Tedore, Jan Funke, Stanley Heinze

## Abstract

Connectomics has become essential for the study of brain function, yet for most research groups it remains prohibitively costly in imaging time, data storage, and analysis. Here, we present an imaging, processing, and analysis pipeline for multi-resolution image acquisition and circuit reconstruction. Applied to the central complex of six insect species, we were able to obtain global projectomes at cellular resolution (40-50 nm) with embedded local connectomes describing key computational compartments at synaptic resolution (8-12 nm). We provide standardized protocols for volume EM sample preparation, image acquisition and image alignment, combined with existing methods for µCT block trimming, automatic segmentation, synapse detection, collaborative skeleton tracing with CATMAID, and segmentation proofreading via CAVE. We validated our workflow by reconstructing head direction cells across all six insect species, which revealed deep conservation at the level of cell types, cell numbers and projection patterns, while also revealing circuit level specializations. Overall, our pipeline democratizes comparative connectomics by making this method accessible for small research groups with modest resources.

## Introduction

Over recent years, studies of brain circuitry have been propelled forward by volume electron microscopy (vEM), the gold standard for producing synaptic resolution maps of neural tissue (connectomes). By generating a structural backbone of the computational architecture of the brain, these connectomes aim to provide an understanding of neural computations at single cell resolution, thereby creating testable hypotheses about the function of neural circuits.

Methods such as serial block-face scanning electron microcopy (SBEM), focused ion-beam SEM, grid-tape EM, and recent powerful variations of these techniques (reviewed in ***Peddie et al., 2022***) have enabled the acquisition of large image volumes of nervous systems at synaptic resolution such as the fruit fly larval nervous system (***Winding et al., 2023***) and adult brain (***Scheffer et al., 2020***; ***Dorkenwald et al., 2024***; ***Bates et al., 2025***; ***Berg et al., 2025***), single columns of the mouse visual cortex (***Bae et al., 2025***), or a 1 mm^3^ cube of the human cortex (***Shapson-Coe et al., 2024***). Using state of the art imaging approaches, the current upper limit of image volumes at synaptic resolution is approximately 1 mm^3^. While the insights from these datasets have been transformative, they are also costly to acquire and take many person-years to fully analyze, thus making high throughput generation of connectomics data challenging. These challenges are particularly evident for large brains and thus prevent us from employing connectomics for large-scale comparative studies or detailed descriptions of inter-individual variations of neural circuitry. Such studies are needed, however, to reveal mechanisms underlying circuit evolution, individuality, or defining ancestral blueprints of fundamental brain circuits (***Barsotti et al., 2021***).

Importantly, methods such as automatic segmentation and synapse detection have leveraged machine learning to drastically accelerate neuron reconstruction (***Buhmann et al., 2021***; ***Sheridan et al., 2023***) and extraction of neural circuits. In particular, collaborative, cloud-based platforms such as CAVE (***Dorkenwald et al., 2025***) have created the possibility to parallelize the manual correction of the raw output from machine learning based neuron segmentation. Compressing 33 person-years into a much shorter time, this platform enabled the generation of a fully annotated, whole brain connectome in the fruit fly (***Dorkenwald et al., 2024***). The same platform was subsequently used to generate a whole central nervous system connectome of the fly (***Bates et al., 2025***) and is currently employed to proofread mammalian cortex segmentation data (***Bae et al., 2025***). Nevertheless, even with these computational advances, the reconstruction of the FlyWire fruit fly brain took approximately five years of proofreading, carried out by hundreds of scientists and many more citizen scientists, to yield a fully annotated connectome (***Dorkenwald et al., 2024***). This demonstrates that further technological advances are required until we can generate brain connectomes routinely, similar to how we can generate whole genome sequences routinely today. Whereas these advances will likely happen over the next decade, these methods will likely remain costly and restricted to be used by larger consortia backed by significant funding. Thus, the aim of our study was to fuse a range of current technologies into a novel, standardized pipeline and thereby enable the acquisition of neural circuit maps at modest effort and comparably low cost. We believe that this approach will aid the democratization of connectomics and thus directly facilitate comparative connectomics across many species and individuals.

One important insight from the connectomes of the fruit fly brain was that, in selected brain regions, many circuit level computations can be approximated without synaptic resolution data. This is due to the tight structure-function relation found in the regular neuroarchitecture of those regions. In these cases, the projectome (***Kasthuri and Lichtman, 2007***), i.e. an exhaustive map of neuronal projections, can predict neural computations to a surprising degree (***Sayre et al., 2021***; ***Pisokas et al., 2020***; ***Turner-Evans et al., 2017***). This is notably the case for the central complex (CX), a region of the insect brain in which projection patterns are synonymous with vector operations essential for insect navigation (***Hulse et al., 2021***; ***Heinze, 2024***; ***Honkanen et al., 2019***; ***Gillet et al., 2025***). Obtaining these insights using projectome level reconstruction relaxes the demands on image resolution compared to what is needed for a connectome, and could in principle be achieved using X-ray imaging even for large volumes (***Kuan et al., 2020***). Due to the coarser image resolution, a projectome also produces less data, and therefore incurs much lower acquisition cost and analysis time than a connectome. Yet, projectomes merely predict connectivity, leaving unanswered whether these predictions are indeed met by synaptic connectivity in all circuits studied.

In this paper we describe an approach in which we combine the advantages of the projectome with those of connectomes by fusing both into a multi-resolution image acquisition and processing strategy. Using this novel data acquisition and processing pipeline we have obtained and analyzed volume EM image stacks of the central complex of a sweat bee, an army ant, a locust, a praying mantis, a cockroach, and an earwig. In short, we used block-face scanning electron microscopy (SBEM) to image the central complex at two resolutions: cellular resolution images (40-50 nm) provide an overview of the entire region of interest, while key compartments are tiled with overlapping synaptic resolution images (8-12 nm). After image acquisition, we use SOFIMA (***Januszewski et al., 2024***) to align images at both resolutions into a coherent frame of reference, allowing for seamless transition between cellular and synaptic resolution reconstructions. Gross neuron backbone morphology is extracted by manual tracing using CATMAID (***Saalfeld et al., 2009***), while fine branches in synaptic resolution compartments are automatically segmented using machine learning (***Sheridan et al., 2023***). Neuron segmentation is collaboratively proofread using CAVE (***Dorkenwald et al., 2025***). Finally, we assign synapses to identified neuron types using automatic synapse detection with machine learning (***Buhmann et al., 2021***). The regularity of the central complex allows us then to combine the global projectome with the local connectomes to extrapolate the connectivity principles of the entire central-complex circuit, comparatively across species. We estimate that our multi-resolution image acquisition strategy was on average 4.5 times faster than if the same region was imaged fully at synaptic resolution, reduced data volumes by approximately the same factor, and reduced the demand on proofreading to a small fraction of comparable full resolution data. Overall, our protocol is employable with modest computational resources, limited personnel and funding typically available to small research groups, thus aiding the general application of connectomics approaches to target problems of comparative neuroscience.

## Results

### A multi-resolution SBEM imaging strategy

In this study we developed a novel approach for mapping neural circuits in brain regions consisting of repeating computational units. Our original motivation for developing this protocol was to acquire connectomics data of an insect brain region called the central complex, from multiple species across the insect phylogeny. These connectomes aimed at describing the central-complex circuits and investigate their evolution. Anatomically, this highly conserved brain region comprises a collection of neuropils that, together, are involved in navigational decision-making (***Gillet et al., 2025***; ***Heinze, 2024***; ***Pfeiffer, 2023***; ***Wilson, 2023***; ***Hulse et al., 2021***; ***Honkanen et al., 2019***). Neurons in the central complex establish repeated vertical columns that function as an array of computational units. These vertical columns are additionally intersected by orthogonally arranged neuronal layers. This architecture is key to the function of the central complex and is already evident at the level of neural projection patterns.

We leveraged the regular neuroarchitecture of our target region to devise a multi-resolution imaging approach using serial block-face electron microscopy (SBEM, ***Denk and Horstmann, 2004***; method also used in ***Sayre et al., 2021***). We used a Thermo Fisher VolumeScope equipped with a VolumeScope directional back-scatter detector (VS-DBS; ThermoFisher Scientific) for image acquisition under low vacuum conditions, with tiling planned using the MAPs 3 software by Thermo Fisher Scientific. However, any comparable microscope with the capability to acquire multiple image tiles at different resolutions would be suitable for our method. In each species imaged, we first scanned the entire central complex at cellular resolution (40-50 nm; Figure 1d-e), aimed at reconstructing all neuron backbones to obtain an inventory of cell types and their projection patterns (Figure 4a). Simultaneously, from the same sample, several sub-volumes were imaged at synaptic resolution (8-12 nm; Figure 1f). As each sample section was imaged at both resolutions, the smaller synaptic resolution areas fully overlapped with the cellular resolution overview image (Figure 1d-e). At both imaging levels, we optimized the voxel sizes for each species independently, aiming at minimizing imaging time while achieving a resolution sufficient to resolve all structures of interest: The backbones of all neurons in the overview data, and synapses and fine dendrites in the synaptic resolution images.

**Figure 1.**
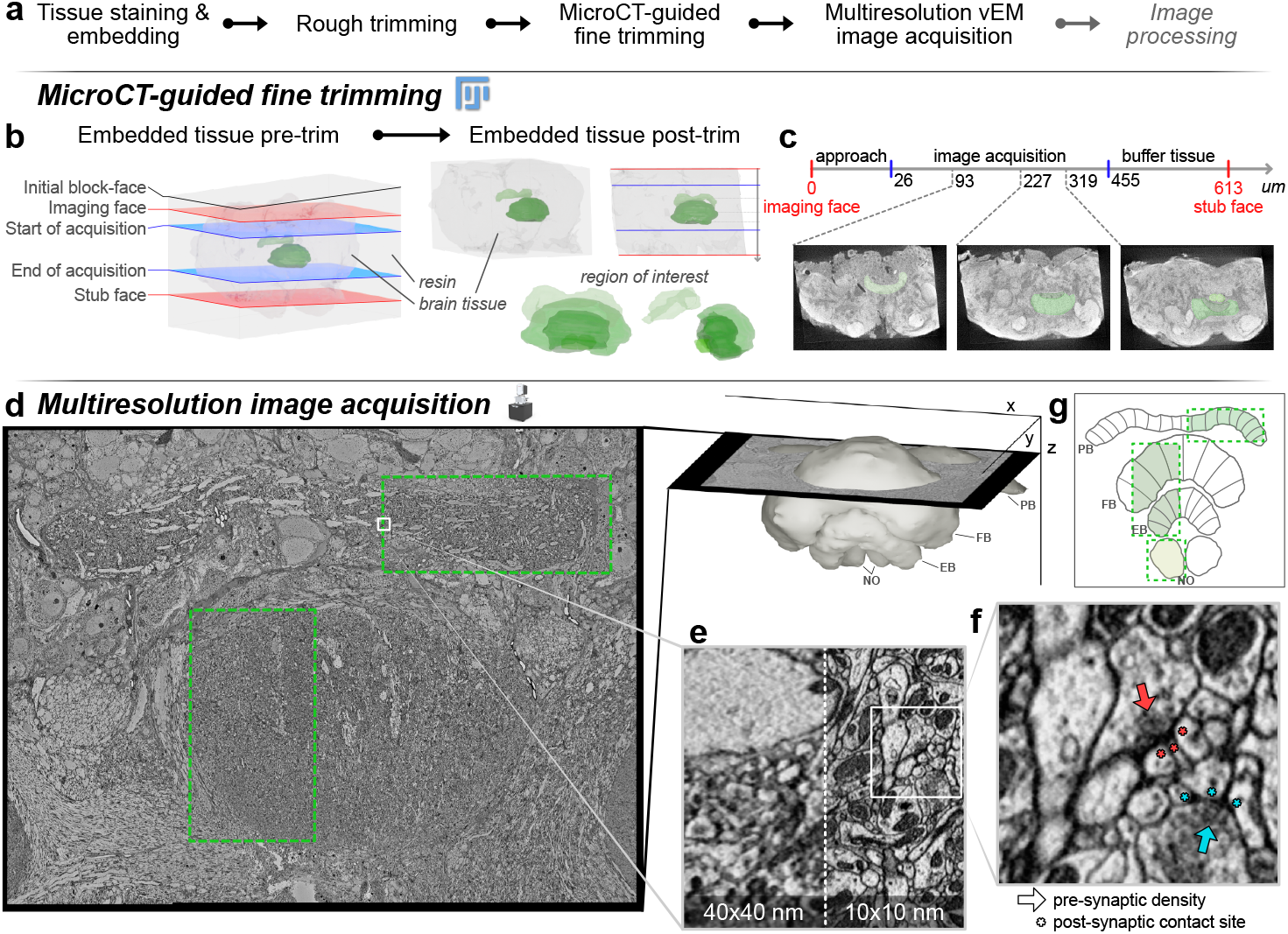
Image acquisition workflow. **a**. Flowchart of the steps needed to acquire images. µCT-guided trimming is optional but facilitates trimming and image acquisition. Shown are examples from the desert locust **b**. Micro-CT guided trimming is done using the FIJI plugin Crosshair (***Meechan et al., 2022***). The imaging face and stub face are defined manually and Crosshair provides the ultramicrotome parameters to trim the original block (left) into the correct orientation (right) as defined by the imaging and stub face planes. Example data comes from the locust brain. **c**. Hypothetical timeline of the sample in b, showing milestones as they can be found at variable depths in the block. **d**. Example of a cellular resolution overview image of the sweat bee central complex, with synaptic resolution images perfectly overlapping (green rectangles). The depth relative to the central complex is shown on the right. **e**. Side-by-side example of cellular resolution (40x40 nm) and synaptic resolution (10x10 nm) images showing the same region. **f**. Zoomed in image showing an example of two polyadic synapses (with multiple downstream partners). Arrows point to pre-synaptic density, asterisks of the same color show corresponding post-synaptic sites. **g**. Schematic overview of the multi-resolution imaging approach. The central complex is tiled by repeating columnar units across its left-right axis. Synaptic resolution tiles (green) are placed such that they follow the path of key computational units covering about half of the central complex. **Figure 1—figure supplement 1**. Comparison between cellular and synaptic resolution image data.

The synaptic resolution image volumes were placed along the path of neurons belonging to the same computational unit (Figure 1g and 3f), thus covering full neural morphologies even if these cells traversed several of the central-complex neuropils. Within these compartments we extracted neural morphologies at the synaptic level to create local connectomes. With this strategy, we effectively constructed projectomes of each insect species’ entire central complex, with nested local connectomes describing a representative set of its computational units (***Kasthuri and Lichtman, 2007***; ***Hildebrand et al., 2017***; ***Sayre, 2022***). Both levels of data were then combined to extrapolate connectivity motifs across the entire central complex, i.e. predict connectivity also for regions only covered by cellular resolution image data. Importantly, by exploiting *a priori* knowledge of the anatomical layout of the region of interest, our method drastically reduces imaging time, computational demands, analysis time and, therefore, the cost of connectomics data acquisition.

As proof of concept, we obtained connectomics data from six large insect species, sampled across the insect phylogeny. This became feasible only as we were able to image at a rate that exceeded that of full synaptic resolution imaging. We estimated the difference between the two approaches and found that our strategy saved a total of at least 362 days of acquisition over the three datasets considered for these estimates, equivalent to 4.5 times faster imaging on average (see Table 1). Although more difficult to quantify, additional substantial time gains were enabled by the much smaller data volumes, significantly accelerating image alignment, segmentation, proof-reading and annotation. Overall, these time gains, and the directly associated lower need for computational resources and personnel, enabled pioneering comparative connectomics work to be carried out in a small research group.

**Table 1.**
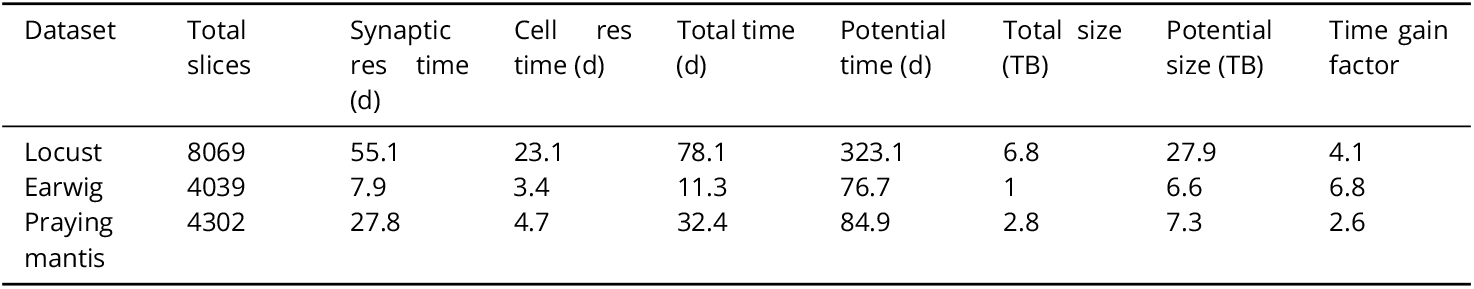
Estimated time gains from multi-resolution imaging approach. Size estimation assumes raw uncompressed greyscale 8 bit image data.

| Dataset | Total slices | Synaptic res time (d) | Cell res time (d) | Total time (d) | Potential time (d) | Total size (TB) | Potential size (TB) | Time gain factor |
| --- | --- | --- | --- | --- | --- | --- | --- | --- |
| Locust | 8069 | 55.1 | 23.1 | 78.1 | 323.1 | 6.8 | 27.9 | 4.1 |
| Earwig | 4039 | 7.9 | 3.4 | 11.3 | 76.7 | 1 | 6.6 | 6.8 |
| Praying mantis | 4302 | 27.8 | 4.7 | 32.4 | 84.9 | 2.8 | 7.3 | 2.6 |

Our vision is that our approach will enable other small research groups to also carry out comparative connectomics work, and thereby provide the wider neuroscience community with the capability to answer research questions that require comparing neural circuits across species, across development, or across individuals. As the outlined imaging strategy lies at the core of our method, it defines the constraints for sample preparation as well as the image processing and analysis pipeline. In the following we will therefore delineate the detailed steps upstream and downstream of the described data acquisition strategy, from sample preparation to connectivity graph analysis, provide the software tools needed to carry out these tasks, and discuss the risks and pitfalls inherent in each step.

### A standardized strategy for sample preparation and quality control

We prepared all samples using a protocol for electron microscopy, originally developed by ***Hua et al***. (***2015***) and adapted in ***Sayre et al***. (***2021***). We further adapted this protocol in two ways. Firstly, we reduced the fixative concentration to one third of the standard concentrations (0.75% Glutaraldehyde, 0.75% paraformaldehyde in 0.1M Cacodylate buffer). Comparing this low concentration fixative side by side with the original recipe consistently yielded superior tissue preservation across all tested insect species. We speculate that the lower concentration fixative more evenly spreads throughout the tissue, particularly in large samples, as no densely fixated tissue layer in the sample periphery is created that might, during high concentration fixation, act as a barrier for effective fixative penetration into the sample center. Secondly, as our region of interest lies at the center of the insect brain and occupies only a fraction of its total volume, we removed peripheral brain regions (optic lobes, subesophageal ganglion) during dissection to reduce sample volume and create large openings in the neural sheath surrounding the brain. Both aid effective fixative penetration. After fixation, samples were infiltrated with heavy metals for contrast enhancement (***Hua et al., 2015***; ***Watson, 1958***) and embedded in a resin block. These blocks were manually trimmed into a cube containing all remaining sample tissue (Figure 1b).

At this step, the sample tissue is dark and opaque, making the recognition of external landmarks close to impossible. To identify the location of the region of interest, we scanned the block using micro-computed tomography (µCT) at resolutions typically between 1 and 2.5 µm (Figure 1b-c). µCT produces 3-dimensional image stacks highlighting regions of high density in solid materials. As the central-complex neuropils are densely packed with heavy metal stained neural membranes, they stand out in a µCT image volume, resembling classic neuropil staining using anti-synapsin immunohistochemistry (Figure 1c). These image volumes allowed us to identify the location of the central complex with a margin of error approximately equal to the resolution of the scan. Besides enabling the identification of the region of interest for SBEM, the µCT data was used to assess the quality of each sample. Any crack, tissue distortion, uneven penetration of heavy metals and other irregularities were visible at this stage and allowed us to select the best specimen of each batch.

In the next steps, the µCT image stacks were further used to finalize the preparation of the sample block for SBEM, and to guide the image acquisition process. Using the µCT scan for guidance, we trimmed the block to a size of maximum 1 mm^3^, matching the size limit for SBEM acquisition and re-orienting it for optimal imaging conditions. For this, we used an open-source FIJI plugin called Crosshair (***Meechan et al., 2022***). This tool provides the angles and distances for positioning the knife of an ultramicrotome by computing the transform between the current block orientation and the desired one, as defined by user input (Figure 1b). The desired cutting plane as well as the anticipated starting position of the SBEM scan were determined based on the µCT image volume.

During Crosshair-guided trimming, the sample was glued to a resin stub with the side of the block facing upwards that would ultimately become the base of the block during SBEM imaging (the stub face). We now used the ultramicrotome to first orient and trim this stub-face surface (Figure 1b). The result of this step was a cleanly trimmed block surface that would constitute the base of the block and was parallel to the anticipated imaging plane. This stub-face surface was now re-glued to a metallic SBEM stub in its final orientation. Next, the newly exposed block face was trimmed by cutting with the ultramicrotome, yielding a block face that was now parallel to the stub face and corresponded to the desired imaging face. The distance to which we trimmed the sample in this way was determined from the µCT data and resulted in a safe starting position for SBEM imaging as well as an imaging plane that was oriented for optimally sectioning through the region of interest.

Two aspects were key during the trimming process. First, we aimed to have as much area containing tissue against the metallic stub as possible, to provide a path of least resistance for electrons to exit the sample during scanning (***Suga and Hirabayashi, 2025***). This essentially grounds the tissue to limit charging during image acquisition, a phenomenon leading to bleaching and damage when electrical charges accumulate in the sample. Second, it was important to place the final block face at a depth far enough from the start of the region of interest to keep space for safe approach, but close enough to limit the number of cuts needed for traversing tissue between the block face and the start of the target volume (Figure 1c).

We found that the optimal orientation of the block significantly facilitated image acquisition by aligning our region of interest with the cutting plane, thereby reducing the need for repositioning imaging tiles throughout scanning. Orienting the block correctly also guided the placement of imaging tiles by aligning identifiable anatomical landmarks along the cutting path towards the region of interest. In the same spirit, one could orient the cutting axis perpendicularly to the shortest side of the bounding box of the imaging field, thus reducing cutting time and tissue loss between slices.

Overall, exploiting µCT imaging for quality control and to guide block trimming via Crosshair was essential to minimize SBEM scan time and to ensure that all regions of interest were safely captured, despite being located within complex, yet completely opaque samples.

### A ready to use pipeline for image alignment

During image acquisition, SBEM produced tens of thousands of images. For the large fields of view used in our imaging approach, the scanning process required the imaged area to be tiled into a mosaic of several individual images per sectioning plane. These partially overlapping tiles must be stitched into one coherent image per slice, before being aligned along the third dimension to create seamless transitions between slices. Our multi-resolution imaging approach additionally produces multiple disparate image stacks of varying resolution, shape, and location. As the various synaptic resolution image stacks only rarely overlap with each other, they must be aligned to the low resolution stack rather than to images of the same resolution (Figure 1d, Figure 2j). Given the special requirements resulting from our imaging approach, we designed a python based, custom alignment pipeline (Figure 2). To this aim, we utilized OpenCV (***Bradski, 2000***) and SOFIMA (***Januszewski et al., 2024***) to compute affine and elastic transformations to align images within their overlapping areas, compensating for offsets as well as local distortions at the image edges. As a strategy, we first aligned all overview resolution images into one coherent image stack (Figure 2b-h), and secondly, used this final overview image stack as reference frame for aligning all synaptic resolution images (Figure 2h-j).

**Figure 2.**
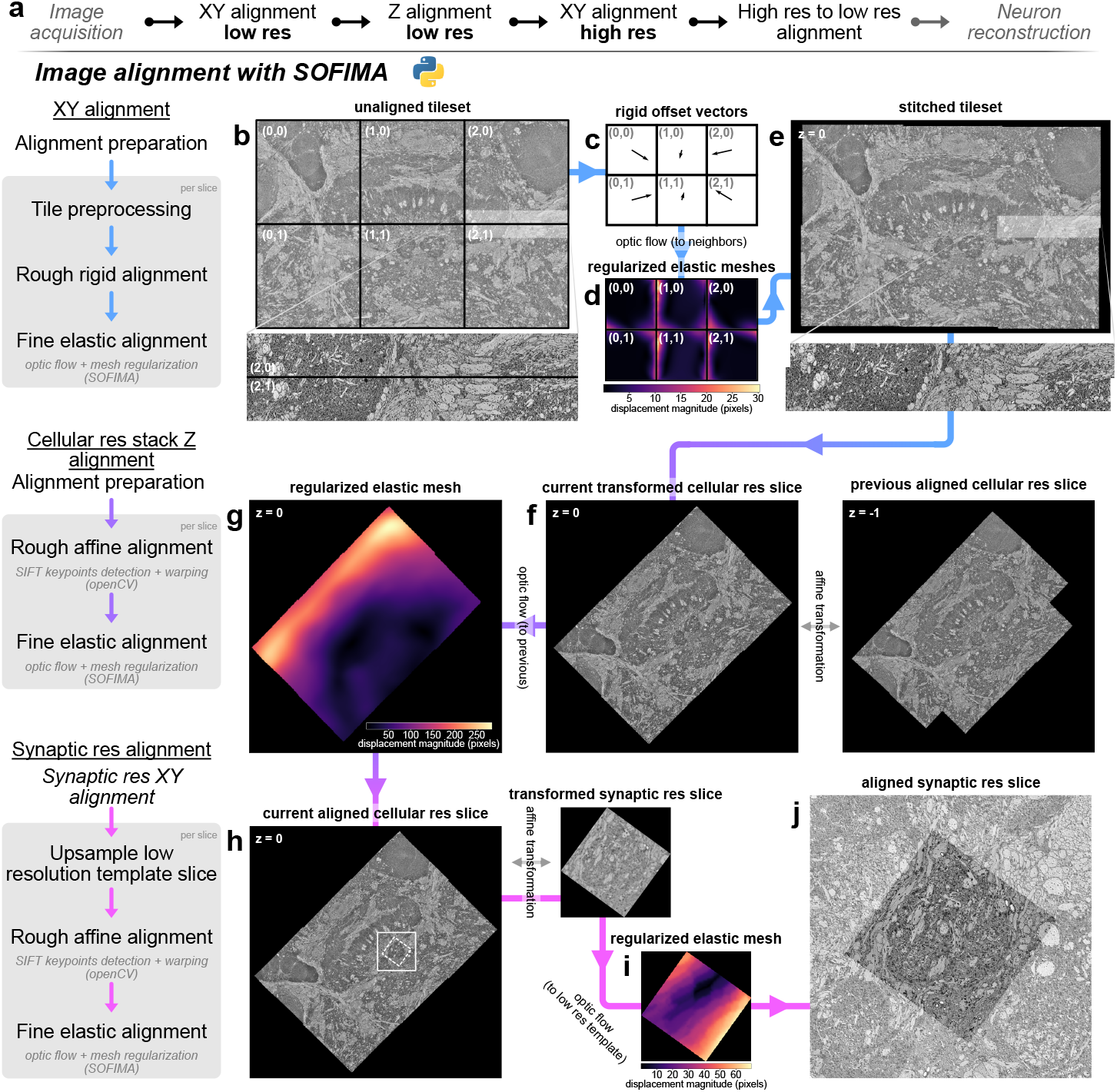
vEM image alignment pipeline. **a**. Flowchart of general steps of the alignment pipeline, shown in detail below. Images shown were acquired from the locust central complex. For large region of interests exceeding the maximum size of a microscope’s field of view, multiple image tiles are required to be assembled into a tileset (**b**). These tiles have overlap necessary to find correspondences with their neighbors. To align them, a rigid offset is first computed using SOFIMA (**c**), before computing fine transformations with optic flow, regularized with an elastic mesh (**d**). The result is one contiguous image per slice with seamless transitions between tiles (**e**). Z alignment is then performed on stitched images by comparing them to neighboring slices along the Z axis. An affine transformation roughly aligns images (**f**) so that fine transformations can be computed using SOFIMA’s optic flow and mesh regularization (**g**). Once the entire cellular resolution overview stack is aligned along the Z axis into one coherent image stack, synaptic resolution images are first aligned in 2D, before being aligned to their corresponding slice of the cellular resolution stack. To this end, an affine transformation is computed to roughly align synaptic resolution to cellular resolution images (**h**) before fine alignment with SOFIMA’s optic flow and mesh regularization (**i**). The final output of this alignment pipeline is one cellular resolution image stack acting as a frame of reference for multiple, perfectly overlapping, synaptic resolution image stacks (**j**). **Figure 2—figure supplement 1**. Alignment of synaptic resolution data to cellular resolution; zoomed in panel j. **Figure 2—figure supplement 2**. Example images showing charging artifacts.

For 2D alignment of the overview image data within the same slice, the coarse placement of image tiles was either inferred by the file metadata and directory tree, or was determined automatically by finding overlap between tiles using keypoint matching with SIFT (scale-invariant feature transform; ***Lowe***, ***1999***). Metadata and file structure are determined by the software used for SBEM image acquisition. While currently written to handle output from the MAPS 3 software (Thermo Fischer Scientific), our pipeline can easily be adapted for different imaging software and microscopes. After coarse placement, image tiles were stitched into one image per slice by computing optic flow, which provides a measure of local displacement. Transformations obtained via optic flow were then optimized with elastic mesh relaxation across each slice separately (Figure 2d). These 2D alignment steps were the same for both cellular and synaptic resolution images.

For alignment of cellular resolution images along the Z axis, each previously aligned slice is compared to its direct neighbor (Figure 2f). Optic flow is computed between neighboring slices, and an elastic mesh is constructed and optimized to produce smooth and minimal deformation (Figure 2g). In some cases, not all stitched images present on a single slice overlapped with each other. Such disconnected image tiles can result from our aim of restricting imaging only to areas of interest within a larger bounding box. This could be the case, for example, when regions on each side of the midline are imaged, but are separated by non-imaged midline adjacent areas to save scan time. Image tiles can thus represent regions of interest that diverged from each other during the scan or that will converge to one image in later slices. To create a frame of reference for disconnected images, alignment in Z therefore always started with the first contiguous slice, and then advanced in both Z directions.

For synaptic resolution stacks, images were aligned to the corresponding slice in the cellular resolution stack rather than to neighboring slices (Figure 2h-j), ensuring perfect correspondence between high and low resolution images.

The implemented alignment pipeline is notably robust to local artifacts, missing slices, and varying image quality throughout the large imaged volumes. This was particularly important as image data quality varied throughout acquisition runs that stretched over weeks, potentially introducing artifacts which would normally result in local alignment mistakes. Key to this robustness was, firstly, the implementation of optic flow computations by SOFIMA which encourages using real resolution and downsampled images simultaneously to capture both global and local deformations. Secondly, metadata is logged in a database throughout alignment to allow smooth continuation after interruptions and to monitor alignment parameters for troubleshooting. This also facilitates restarting specific steps while iteratively adjusting parameters if necessary.

Overall, we found that our pipeline was able to correctly and automatically align image tiles with overlaps as small as 10 pixels, presenting cutting or charging artifacts, and across gaps caused by missing slices. The parameters for elastic alignment we ultimately converged on transfer well between species, and can be expected to provide a solid starting point for any new datasets acquired with our imaging strategy. Our alignment pipeline therefore provides an off-the-shelf tool for transforming raw image output from a SBEM microscope into a fully aligned image stack ready for analysis.

### A two-level neuron reconstruction strategy for connectomics

After generating fully aligned image stacks at cellular and synaptic resolutions, we were faced with the challenge of generating coherent neural morphologies from these data. We followed two complementary reconstruction strategies to extract maximal information from both sets of images. First, cellular resolution overview image stacks were used to manually trace neural backbones across the entire central complex, generating a global projectome. Second, automated image segmentation combined with manual proofreading was used to extract comprehensive morphologies of all neural branches within synaptic resolution image volumes, thus creating the dense reconstructions needed to generate local connectomes (Figure 3e).

**Figure 3.**
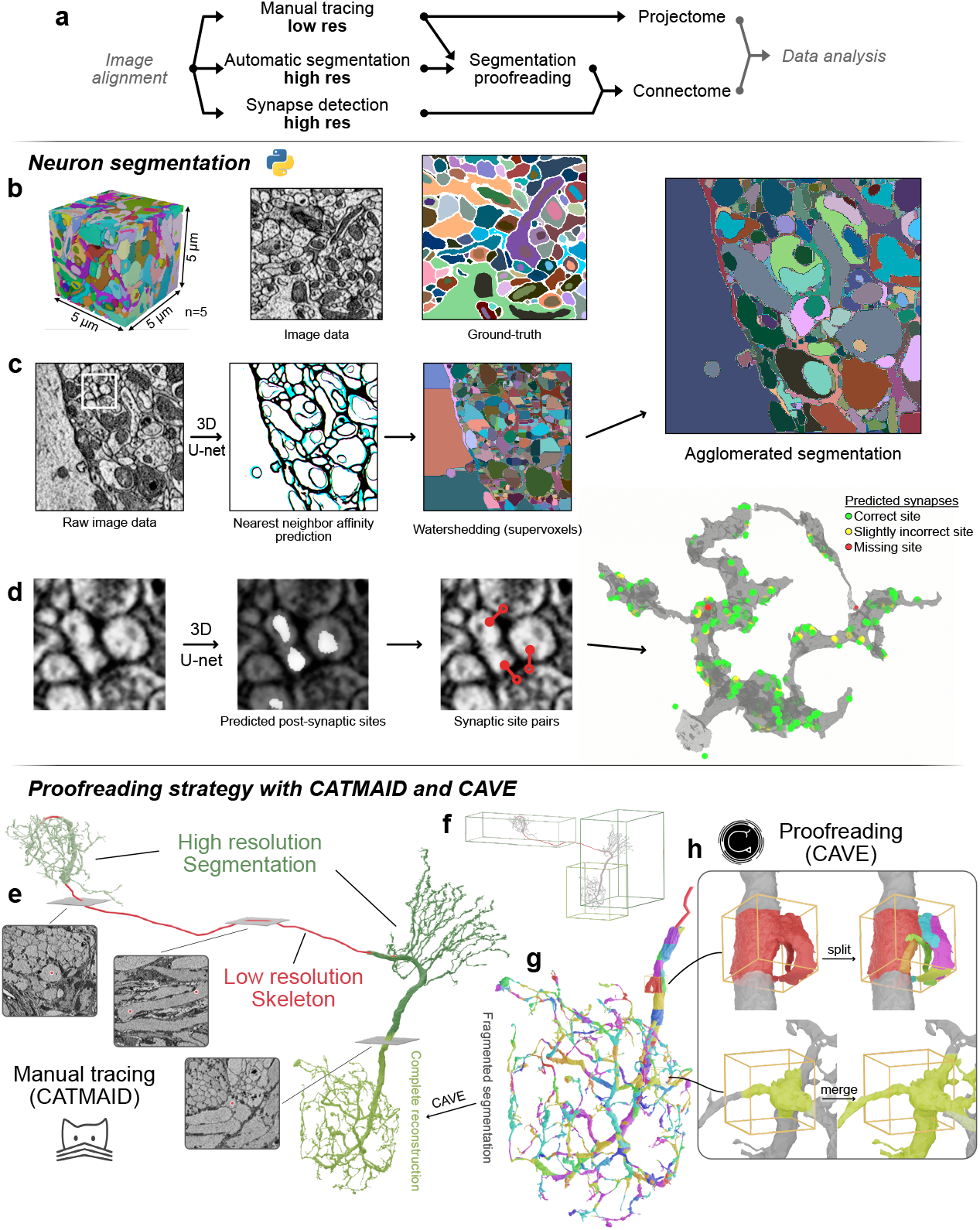
Combining manual and automatic neuron reconstruction. **a**. Flowchart for multi-resolution reconstruction of neuron morphology **b**. Example of ground-truth data manually segmented to train a 3D U-net. We used 5 cubes representing 5x5x5µm^3^ of image data to train our model. **c**. Segmentation workflow. Raw data is used to predict nearest neighbor affinity (NNA) with a trained 3D-Unet. NNA are used to compute supervoxels by watersheding. The agglomerated segmentation consists in relabeling connected components after thresholding of a segmentation graph, producing accurate neuron segmentation. **d**. Example of a synapse (zoomed in from raw image data in c) identified by a 3D-Unet trained to detect synaptic sites. Pre-synaptic (not shown) and post-synaptic sites are detected separately and used to compute synaptic site pairs. The right-hand side shows an example of a reconstructed and proofread neuron with detected synaptic sites. Green: accurately predicted partner and synaptic site. Yellow: accurately predicted partner with wrong synaptic site. Red: false negative, missing prediction. **e**. Overview of a multi-resolution neuron reconstruction. The neuron backbone is manually traced using catmaid (red), while synaptic resolution compartments are automatically segmented (green). **f**. The skeleton resulting from manual tracing is used to identify cell types and bridge the gap between compartments of the same neuron segmented in synaptic resolution image stacks. **g-h**. Illustration of proofreading using CAVE. Segmentation is fragmented into regular chunks of equal size when uploaded to CAVE (g). An existing skeleton can be used to find fragments belonging to the same skeleton, and proofreaders otherwise assemble fragments before merging them. Using CAVE (h), proofreaders can correct split errors caused by faulty segmentation, and merge errors either caused by faulty segmentation or artificial splits. **Figure 3—figure supplement 1**. F-scores for synapse predictions. **Figure 3—figure supplement 2**. Example of fused membrane artifacts.

**Figure 4.**
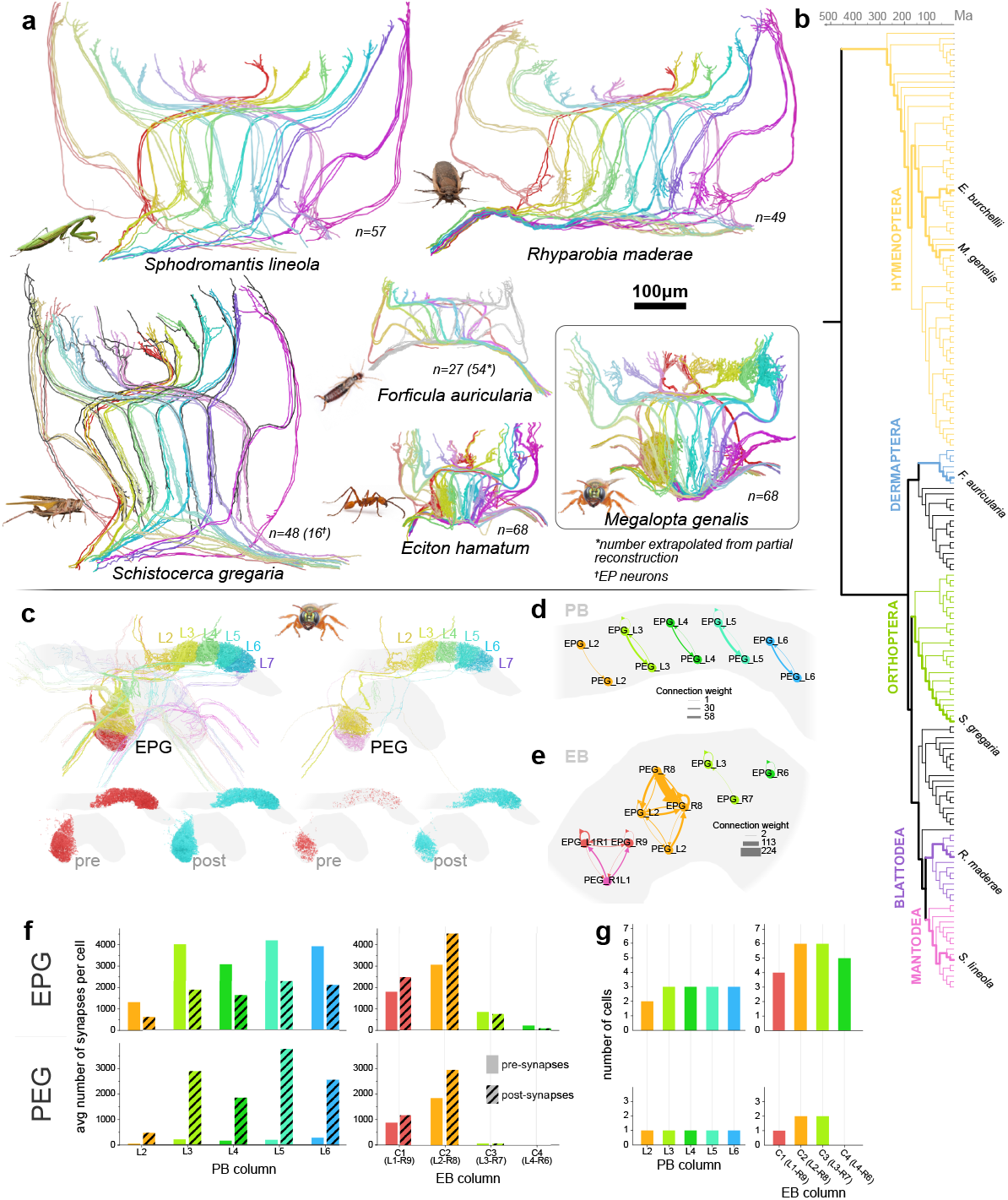
Proof of concept reconstruction of head direction cells across insects. **a**. Traced EPG and PEG neurons, color-coded by the column of the protocerebral bridge they innervate. Insects: African praying mantis (*Sphodromantis lineola*), Madeira cockroach (*Rhyparobia maderae*), desert locust (*Schistocerca gregaria*), European earwig (*Forficula auricularia*), army ant (*Eciton hamatum*), sweat bee (*Megalopta genalis*). For the locust, EPG neurons matched the projection patterns in other insects (n=48), while some EPG-like neurons did not project outside of the central complex (EP neurons, n=16)(shown in dark grey). For the earwig, only right hemisphere cells traced (shown in color). Total cell count (54) was estimated by doubling cell number from the right hemisphere (27), assuming symmetry between right and left side (mirrored right neurons shown in grey). **b**. Phylogenetic tree highlighting the species shown in panel a. **c**. 3D representation of segmented EPG and PEG neurons of the sweat bee. Skeletons bridge the PB and the EB where automatic segmentation and proofreading yielded full reconstruction of neural branches in six columns of the protocerebral bridge and four columns of the ellipsoid body. Automatically detected synapses are shown below for each cell type. Presynaptic sites, red; postsynaptic sites, blue. **d-e**. Connectivity graphs showing recurrent circuits between EPG and PEG cells in the protocerebral bridge and the ellipsoid body of the sweat bee. Skeletons were used to match reconstructions across compartments, enabling detection of recurrent circuits across distant high-resolution image volumes. **f-g**. Synapse distribution (f) and number of cells (g) per column across neuropils for EPG (upper row) and PEG neurons (lower row) for the sweat bee. Note that low number of synapses in columns L7 of the protocerebral bridge, and C3 and C4 of the ellipsoid body are due to incomplete reconstructions at the boundary of synaptic resolution image stacks. Picture credits in panel a: Chris Dlouhy (*Rhyparobia maderae*), Daniel Kronauer (*Eciton hamatum*), and Ajay Narendra (*Megalopta genalis*). Picture of *Forficula auricularia* by Georg Ekhöft, Observation.org (https://observation.org/photos/109145418/), CC BY-NC 4.0 (https://creativecommons.org/licenses/by-nc-nd/4.0). **Figure 4—figure supplement 1**. Side-by-side comparison of EPG and PEG neuron in the protocerebral bridge.

We used the collaborative annotation tool CATMAID (***Saalfeld et al., 2009***) for manual reconstructions of projectome level data. While cellular resolution images were exclusively reconstructed with this method, we also applied manual tracing to explore synaptic resolution image volumes when needed. As outlined below, these manually traced neural branches provide important guidance for proofreading of automatically segmented data (Figure 3e, Figure 4a,c). CATMAID is an efficient, open-source software that has been widely used for collaboratively reconstructing neural skeletons across insect brain datasets in *Drosophila*. Limiting reconstructions to skeletons, i.e. minimal representations of a neuron’s morphology, captures the most essential defining features of cell types in our datasets, often within minutes for an individual neuron. This speed was possible as we only traced the backbone of a neuron and their major, type-defining branches. As a result, we obtained neuropil-wide projection patterns of anatomically identified major cell types of the central complex of many insect species with comparably little effort. At the same time, these neural backbones interconnected all synaptic resolution image volumes within any individual species, thus attaching cell type identities to branches that were segmented from these higher resolution data (Figure 3f).

Fine branches and synapses are required to build local connectomes, but can reliably only be identified within synaptic resolution image volumes (Figure 3e, 4c). While, in principle also traceable with CATMAID, manual tracing of extended arborizations of very fine fibers of many hundreds of neurons is very time consuming. Additionally, the terminal morphology of neurons often was too complex to be effectively captured by a minimalistic, skeletonized backbone. To overcome both limitations, we trained a convolutional neural network (CNN) to predict nearest neighbor affinity and local shape descriptors (***Sheridan et al., 2023***) to segment all neuronal profiles within our synaptic resolution data (Figure 3c). Our training and prediction pipelines follow the method published by ***Sheridan et al***. (***2023***) and are written in python. We implemented the gunpowder (https://github.com/funkelab/gunpowder/tree/main) pipeline for data augmentation and the package daisy (***Nguyen et al., 2022***) for parallel processing.

Our training datasets were obtained from the tropical bee *Megalopta genalis* and comprised four 5x5x5 µm^3^ image cubes from three different synaptic resolution image stacks. Within these cubes we manually segmented all neurons as ground truth for training our model (see Methods for details on the training procedure). The result of this training was a surprisingly robust model that could be readily applied across species, brain regions, and different imaging methods. While possessing some limitations, our model is a ready-to-use starting point for segmenting a broad range of volume electron microscopic data.

As for any volume electron microscopical datasets, the quality of our image data varied during acquisition. This produced localized artifacts such as tissue damage, charging, or misalignment, causing two types of errors in the segmentation: false splits (neurons incorrectly separated) and false merges (neurons incorrectly fused; Figure 3g). To correct these errors and thus generate complete neural branching trees, manual edits were required. To this aim we deployed the CAVE platform (***Dorkenwald et al., 2025***) for collaborative proofreading of neuron segmentation.

Taking advantage of the two level image data generated by our acquisition strategy, our proof-reading pipeline combines CATMAID and CAVE to produce hybrid representations of neuron morphology (Figure 3e). In practice, proofreading is initiated by tracing all backbones of neurons of interest until reaching first or second order branching points. As a next step the segmented fragments that belong to these neurons are identified by overlapping the skeletons with the raw segmentation output. To facilitate this process, we developed an application deployable alongside CAVE which allows users to query skeletons from a CATMAID instance and identify the corresponding fragments in a CAVE volume. This provides an advanced starting point for proofreading even in highly fragmented segmentation data. The final proofreading process then comprises the identification of any missing neural fragments, correction of any merge errors (Figure 2), and fusing all fragments into one contiguous and complete neural branch.

Once proofreading was completed for each synaptic resolution branch of a neuron of interest, these branches are combined with the CATMAID based skeleton into a full neural morphology.

### Combining multi-resolution morphologies with synapse predictions to extract neuronal circuits

To extract connectomics data from our segmented image stacks, proofread neurons need to be interconnected via synapses. As manual identification of synapses is extremely time-consuming in large datasets, we employed machine learning for automatic synapse prediction. We used a model architecture and method published by ***Buhmann et al***. (***2021***) to train a CNN to identify pre- and postsynaptic site pairs in our synaptic resolution image data (Figure 1f). We were able to achieve results comparable to the original study when validating our model on image data from the sweat bee’s neuropils, despite image quality varying significantly between image regions. We calculated F-scores for variable image quality, used as a measure of prediction confidence, of 0.63, 0.71, and 0.52 for best image quality in the protocerebral bridge, fan-shaped body, and noduli respectively (Figure 1). As each synapse is predicted with a confidence score, we applied thresholding to remove low-confidence predictions. Similarly to the segmentation pipeline, training for synapse prediction uses gunpowder (https://github.com/funkelab/gunpowder/tree/main) for data augmentation and daisy (***Nguyen et al., 2022***) for multiprocessing.

Synaptic sites were predicted as voxel coordinate pairs, and then automatically assigned labels corresponding to the underlying segmentation, which itself was linked to cell type identity via manually traced skeletons. Alternatively, labels can also be assigned directly through the CAVE Annotation and Materialization services (***Dorkenwald et al., 2025***). Synapses connecting the same label were removed, as autapses (self-synapses) are generally considered artifacts in automatic reconstruction of insect brain circuits (***Hulse et al., 2021***). Our heuristic therefore removes connections that map between two sites within the same neuron as false positives.

Circuits can finally be extracted by summing the number of synapses connecting neuron-pairs. To this end, we used common python libraries such as pandas, numpy, and networkx. Connectivity data is generally visualized in graph form (Figure 4d-e) or as matrices, using libraries such as networkx, seaborn, or matplotlib for plotting. As insect neurons connect redundantly, across multiple synapses, we removed any connections weaker than three synapses, following conventions developed in *Drosophila* connectomics (***Hulse et al., 2021***). This likely further removes falsely predicted synapses, as those are unlikely connecting the same partners repeatedly, different from true connections.

Together, our approach combines a range of efficient, existing annotation tools for manual tracing, automatic segmentation, and automatic synapse prediction into a unique analysis pipeline tailored to our multi-resolution imaging strategy. The described workflow produces accurate representations of local brain circuitry on a synaptic level from any aligned volume electron microscopy image stack. Importantly, this pipeline can be readily deployed with modest investment in IT infrastructure and can be effectively managed by small research groups.

### Head direction cells across insects as proof of concept

To verify that our approach can generate meaningful insights from a range of large insect species, we acquired image data of the central complexes of six insect species (Figure 4a): *Sphodromantis lineola* (African praying mantis), *Rhyparobia maderae* (Madeira cockroach), *Schistocerca gregaria* (desert locust), *Forficula auricularia* (European earwig), *Eciton hamatum* (an army ant species), and *Megalopta genalis* (a tropical sweat bee). These species were chosen to represent a wide range of orders within the insect phylogeny. Most of these species are established models in neurobiology, but lack comprehensive descriptions of their head direction circuits. Given their large evolutionary distance of more than 400 million years, as well as vastly different ecologies, we asked, firstly, whether neurons homologous to the fruit fly head direction cells exist across species. In each species, we first created projection level reconstructions of all identifiable EPG and PEG neurons (also called CL1 in non-fly insects), known to be the main head direction cells in the fruit fly *Drosophila* (***Seelig and Jayaraman, 2015***). These cell types were identified as EPG/PEG neurons by their unique anatomical projections described in the fruit fly connectome and by individual dye-injected examples from other insects. As in all reported examples, we found EPG/PEG cells to be columnar neurons that connect specific columns between the protocerebral bridge and the ellipsoid body, and project contralaterally, presumably to the gall (a neuropil external to the central complex, not included in our image volume). Cell types equivalent to fly EPG/PEG and locust CL1 neurons have therefore likely emerged early in the insect phylogeny.

While individual examples of EPG-like neurons have been reported from many insect species before, we were able to describe the complete repertoire of these cells for the first time. As in flies, we consistently identified four individual EPG/PEG cell types in each regular column of the central complex in all holometabolous species. In hemimetabolous species (locust, cockroach, praying mantis, earwig) this set was reduced by one cell, so that three neurons innervated each column. Consistently, cells originating from the midline adjacent columns of the protocerebral bridge contained fewer cells. In the locust, a fourth EPG-like cell was present in each column, but lacked the axonal projection towards the gall and was therefore not included in the cell count.

As the projection patterns of EPG neurons in *Drosophila* provides the backbone for the computations underlying head direction coding, we next asked whether these projection patterns are conserved. Across all species, EPG/PEG cells tiled the PB into nine columns per hemisphere, named after their position from the midline: L1 and R1 are the closest to the midline for the left and right hemisphere respectively, whereas L9 and R9 are the most lateral columns. Similarly, EPG/PEG cells also tile the EB into nine columns, named C1 to C9 from the right to the left side. These three sets of columns were connected in a stereotypical pattern, which was almost completely identical to that found in flies and bumblebees, and that of CL1 cells originally described for the locust by ***Williams*** (***1975***). In each species, EPG/PEG neurons projected each hemisphere of the protocerebral bridge onto the complete width of the ellipsoid body, so that neural signals are sent from a single ellipsoid body column to two corresponding columns of the protocerebral bridge, one on each hemisphere (Figure 4). While these patterns were highly conserved across most of the central complex, more variability was found at the midline adjacent columns of the protocerebral bridge. Here, EPG/PEG neurons differed in their innervation patterns of the protocerebral bridge and branches were either contralateral to the soma fiber, ipsilateral, or bilateral.

As *Megalopta genalis* was the first image volume acquired, we used it to develop and test our automatic segmentation and synapse prediction pipeline. We used the resulting data to ask whether EPG neurons, PEG neurons and their interconnectivity were conserved at the synaptic level (Figure 4c-g). We thus segmented and proofread all EPG/PEG neurons in the six columns of the protocerebral bridge and in two of the four columns of the ellipsoid body that were covered by our synaptic resolution volumes of the bee central complex (Figure 4c).

While EPG and PEG neurons have identical projection patterns, both types differ in their polarity and, consequently, in the main direction of information flow. This polarity difference has been reported in light microscopy studies, based on beaded versus smooth arborization structure in either the protocerebral bridge or ellipsoid body, and was verified by synapse distribution data based on EM images in *Drosophila*. This observation was confirmed in our *Megalopta* data, albeit with less pronounced polarity difference in the ellipsoid body. Generally, EPG and PEG neurons have mixed terminals in both neuropils, but EPG neurons are predominantly presynaptic in the protocerebral bridge and postsynaptic in the ellipsoid body, while PEG cells have the opposite synapse distribution, with almost purely dendritic terminals in the protocerebral bridge (Figure 4f). These data also confirmed that out of the four EPG-like cells in each column, three are EPG neurons and one belongs to the PEG type, again resembling the configuration of the fly.

While a detailed analysis of the synaptic connections of the bee’s EPG neurons is beyond the scope of this paper, we identified synaptic connections between EPG and PEG neurons as a proof of concept. Those cell types form recurrent loops between the protocerebral bridge and the ellipsoid body, similar to what has been reported for *Drosophila* (Figure 4d-e). In more detail, in the protocerebral bridge, EPG neurons provide strong input to PEG neurons of the same column. As PEG neurons are almost exclusively dendritic in the protocerebral bridge, they do not synapse onto EPG cells at that location. However, in the ellipsoid body, EPG neurons provide input to each other and form balanced, reciprocal connections with PEG cells. Therefore, in bees, the recurrent loop between EPG cells and PEG cells is directly closed, unlike in *Drosophila*, which lacks direct PEG to EPG connections in the ellipsoid body and closes this loop indirectly via PEN neurons (***Turner-Evans et al., 2020***; ***Hulse et al., 2021***). This difference in one of the core feedback loops of the head direction system suggests that despite the universal need to encode heading across insects, this circuit retains a surprising degree of evolvability.

## Discussion

### A robust connectomics workflow

The field of connectomics has rapidly expanded over the last few years, producing groundbreaking datasets describing the nervous systems of invertebrates (***Witvliet et al., 2021***; ***Verasztó et al., 2020***; ***Cook et al., 2019***), such as the brain of fruit flies (***Dorkenwald et al., 2025, 2024***; ***Winding et al., 2023***; ***Scheffer et al., 2020***) or fragments of larger vertebrate brains (***Lee et al., 2016***; ***Shapson-Coe et al., 2024***; ***Bae et al., 2025***; ***Shvedov et al., 2025***). These advances have been driven by large consortia capable of withstanding the high cost of such endeavors. The ability of smaller research groups to perform studies using connectomics methods remains highly limited. Major remaining obstacles to democratizing connectomics currently are the time and cost required for image acquisition and processing, data storage and analysis, as well as the expertise required to train and apply advanced machine learning models. These challenges have been sufficiently discouraging for small research groups with limited resources to employ connectomics methods, even aiming at small image volumes or partial brains. This severely limits the questions that can be asked using these methods and prevents our understanding of synaptic level organization of neural circuits from going beyond the currently mapped model organisms. Many pressing questions requiring answers at the level of neural circuits remain therefore out of reach, including questions aimed at unraveling developmental processes, evolutionary principles, the neural basis of individuality, or structural changes due to learning or pathological conditions.

With the workflow detailed in this paper, we aim at making connectome reconstructions cheaper and more accessible, facilitating imaging a broader range of samples and enabling repeated imaging under different experimental regimes. Firstly, we demonstrated that our multi-resolution approach accelerated imaging compared to a similar volume acquired at synaptic resolution by a factor of 4.5 on average, reducing imaging time from months to weeks. At the same time, the amount of raw data produced was reduced by the same factor, lowering the computational demands on data storage, handling, and processing. As our strategy confines imaging to regions of interest that are selected based on concrete hypotheses, reconstruction efforts are ultimately more targeted compared to conventional connectomics, focusing on key circuits and thereby gaining additional time during analysis of the acquired data.

Secondly, our workflow, from sample preparation to connectivity analysis, can be applied to many types of samples without adaptation. Despite using identical sample preparation protocols, and largely identical imaging setting, we were able to produce high quality image data. Although of somewhat poorer quality than those used for exhaustive full-brain connectomics in *Drosophila*, our data comprised sufficient tissue integrity and contrast to consistently allow for the reconstruction of local connectomes. Being able to apply an identical histology protocol (adapted from ***Hua et al., 2015***) across many insect species is an essential prerequisite for broad, comparative studies. Optimizing protocols for any particular species often requires substantial efforts and time investment and becomes unfeasible when aiming at imaging more than a handful of species.

Although our strategy likely sacrifices optimal image quality for any particular species, flexibly combining manual tracing and automatic segmentation during neuron reconstruction grants sufficient robustness during analysis to describe neural circuits at a synaptic level, across regions of interest spanning hundreds of micrometers. Notably, we have not employed sophisticated image processing strategies to compensate for detriments in image quality. Adding smart image restoration algorithms to our workflow could thus offer the possibility to further increase the robustness of our overall method.

Similarly to our sample preparation and imaging protocols, our freely available segmentation models are readily transferable between species. Automatic segmentation methods using machine learning traditionally require large amounts of ground-truth and computational resources to be efficient and are often outsourced to commercial service providers, substantially increasing the overhead cost of connectomics. In contrast, our segmentation pipeline, utilizing methods developed by the Funke lab (***Sheridan et al., 2023***), was trained with ground-truth data produced from only a single dataset. Yet, it yielded a model capable of segmenting image data of varying quality as well as data from multiple different insect species, covering a substantial size range. This transferability was not limited to central-complex image data acquired under similar conditions (Figure 4), but our model was also successfully applied to data from the insect mushroom body (M. Kinoshita, unpublished data) as well as the optic lobe and retina, all acquired with a different microscope (K. Arikawa, G. Belušič, A. Stöckl, unpublished data).

In summary, our workflow and the associated python pipelines can be readily used to image, process and analyze volume EM data, from start to finish and with little optimization requirements. While optimizing and adapting settings will likely produce better results, using our workflow as an off-the-shelf solution can be expected to at least provide a valuable starting point for a wide range of datasets aimed at producing connectomes, without requiring advanced technical knowledge or large financial investments.

### Applications beyond the insect central complex

Our multi-resolution strategy was developed with the intent of studying the specific neuroarchitecture of the insect central complex, which is characterized by its columnar units repeating in a highly predictable arrangement. In principle, our workflow is therefore applicable to any modular brain region, where combining long-range projections with local connectivity maps yields meaningful insights, i.e. where local, synaptic resolution connectomes can be embedded in a cellular resolution, global projectome to gain insights into overarching computational principles. This readily applies to brain regions with regular neuroarchitecture, such as the insect optic lobes, featuring layers and columns, the mushroom bodies with their repeating output modules (***Li et al., 2020***), or the glomerular organization of the insect antennal lobe. Importantly, repeating columns and layers that are interconnected by long range fibers, as well as glomerular architectures, are not limited to insects, but are also well known from vertebrate brains. Studying small, representative portions of the cerebral or cerebellar cortices of small mammals, the tectum opticum across vertebrates, or brain stem nuclei should be viable applications of our imaging and analysis strategy.

The targeted approach and the associated cost and time efficiency additionally allows studying any of the brain regions mentioned above across developmental stages, between individuals, life stages, after experimental manipulation, or between pathological and healthy conditions. This includes, for instance, the study of circuit level effects of learning, developmental conditions, pesticide exposure, or comparison of migratory versus non migratory stages of long distance migratory species.

In principle, our pipeline is even agnostic to the used imaging modality for cellular resolution data. With advances in X-ray imaging pushing the limits of resolution to nanoscales, combining cellular and synaptic resolution to reconstruct brain circuits can be extended to correlative imaging methods. One could, for example, use high resolution X-ray imaging to acquire a cellular resolution projectome of a sample at state-of-the-art sub-100 nm voxel size (***Kuan et al., 2020***), and subsequently image compartments of the same sample at synaptic resolutions routinely achieved with volume EM. By using these techniques one can envision imaging of much larger projectome volumes, combined with numerous local connectivity maps representing different brain regions. While not reaching the comprehensiveness of complete maps of information flow derived from whole brain connectomes, the insights into computational principles gained from a multi-resolution imaging and analysis approach might be comparable and, importantly, are feasible without massive financial investments.

### Limits of our approach

In our workflow each step relies on the correct execution of the preceding ones, from sample preparation to image segmentation. Yet, it is the quality of the acquired images that ultimately sets the limits for efficient neuron reconstruction, accurate synapse prediction and acceptable proofreading demands.

We built our strategy based on classic SBEM imaging, a tradeoff between availability and reliability against speed. In our approach, the maximal volume that can be imaged is around 1 mm^3^, which is set by the maximal size of the block accepted by the SBEM microscope. This is comparable to the the state-of-the-art maximum volume that can be acquired using other volume EM techniques at full synaptic resolution (***Shapson-Coe et al., 2024***). Going beyond this size limit, e.g. to increase our overview image volumes, would require the use of nano-scale X-ray based imaging, or physically partitioning a larger sample into smaller cubes, e.g. by near lossless hot knife sectioning (***Hayworth et al., 2015***). However, these methods require access to highly specialized equipment not readily available to most research groups, so that, in practice, an upper boundary for imaging volume is set by the used microscope.

Beyond general sample size, maximal acquisition speed is dependent on the propensity for a sample to charge, which defines imaging parameters. Highly conductive samples can sustain higher electron dose without charging, allowing lower dwell times to yield satisfying signal-to-noise ratios and high contrast. With our histological protocol and the used microscope hardware, charging prevented us from imaging at high vacuum conditions, and we therefore employed a low vacuum setting. In this condition, water droplets in the imaging chamber’s atmosphere help charge dissipation (***Suga and Hirabayashi, 2025***). While granting acceptable contrast without sample charging, the required pixel dwell times were long, thus yielding relatively long overall scan times. With this imaging mode, large samples can take weeks to months of continuous imaging, despite the significant time savings as a result of multi-resolution imaging. For example, the largest sample acquired with this method - the locust central complex (volume: 800 x 680 x 403 µm) - required a total of 78 days of scanning time using SBEM (Table 1). In practice, such long acquisition times increase the probability for microscope malfunction, which may interrupt acquisition temporarily or render slices unusable. The results are, for instance, low image overlap due to drifting of image tiles, missing slices caused by vacuum failure, and additional errors due to inevitable human mistakes when monitoring scans over many weeks. While our data processing pipeline was robust to these errors, there are likely faster alternatives to our low vacuum SBEM imaging mode, as long as they are available to a research group. A relatively straight forward upgrade would be the use of a SBEM equipped with a focal charge compensator (***Suga and Hirabayashi, 2025***; ***Deerinck et al., 2018***), a device that blows nitrogen gas on the sample surface to dissipate electrons, or the use of FIB-SEM for smaller image volumes. Both methods limit sample charging and allow for more aggressive imaging parameters. Importantly, the time saving factors due to multi-resolution imaging translate also to alternative, faster imaging methods, and any downstream processing would be identical. Additionally, custom software such as SBEMimage for the Zeiss 3View microscope (***Titze et al., 2018***) are designed to monitor image acquisition, and implement various quality control and automatic focusing features. Such open-source software could increase the robustness of our strategy and the quality of imaging during long runs, by introducing fail-safes against common instrument issues.

While whole brain imaging at synaptic resolution is expensive and generates data amounts that are challenging to handle in practice, the region of interest is clearly defined as the brain boundary. In comparison, our more targeted workflow requires continuous attention from an experienced neuroanatomist to successfully place and update regions of interest during the scanning period. This applies to the cellular resolution overview, but especially to the synaptic resolution sub-volumes. Initially, we performed manual, on-the-fly placement of imaging tiles, resulting in loss of fiber bundles (e.g. in *Megalopta*) or partial loss of neuropil boundaries at the forefront of imaging (e.g. at the dorsal edge of the protocerebral bridge in *Megalopta*). This need was later mitigated by introducing µCT based trimming and scan guidance, providing landmarks for planning placement of tiles before the image acquisition starts. Nevertheless, expert guidance is essential during image acquisition to ensure capturing the correct brain regions without sampling excessive tissue outside of the target region.

Due to the compromise in accepted sample and image quality, very large diameter axons at neuropil boundaries occasionally generated loss of membrane artifacts that required us to generate a comparably fragmented segmentation output as our default starting point for proofreading. This increased the demand for manual edits during proofreading compared to data resulting from highly optimized histology and optimal segmentation models. Our strategy was to mitigate this state of the data by creating effective interactions between manual tracing software, which is more robust to image artifacts, and CAVE based segmentation editing. Our approach thus saves time by not retraining our segmentation models with new ground truth for each sample and not optimizing sample histology for each species, at the cost of increasing the proofreading burden. Whether this approach grants net benefits to a connectomics project will depend on the research question, number of species targeted and how clearly defined the circuits of interest are. Generally, the resulting off-the-shelf protocol and segmentation models are clearly not optimized for any particular sample, but have the advantage that they can be used as they are for any new species and brain region, at acceptable performance and low cost. Employing exploratory TEM on any sample type before embarking on a large SBEM run is advisable to ensure acceptable tissue preservation.

Finally, as our approach deliberately images only a sub-volume of a brain, neurons are cut at the boundary of the scanning volume. While our strategy is to target specific circuits and cover all key elements fully in the imaged volumes, orthogonally intersecting cells, input neurons from regions outside the image volumes, as well as output cells are bound to be only partially included. Thus, the data resulting from our approach benefits greatly from complementary data to gain full neural morphologies of cut neurons, e.g. via single cell dye injections, Golgi stainings, or immunohistochemistry.

### Deep conservation of primary head direction cells across insects

As a proof of concept, we manually traced neurons resembling the fruit fly head direction cells (EPG/PEG neurons) across six insect species. We additionally segmented and proofread a subset of these cells in the tropical sweat bee *Megalopta genalis* at synaptic resolution and performed initial analyses of their interconnections. For the species studied, individual examples of these neurons were previously described as CL1 neurons in the desert locust (***Heinze and Homberg, 2008***) and the Madeira cockroach (***Jahn et al., 2023***). As these studies identified those cells by dye injections, the full projection patterns, the number of cells per central-complex column and their synaptic connections were not directly accessible. In locusts, the entire projection system of EPG-like cells was described in the seminal paper by ***Williams*** (***1975***), using manual tracing across brain slices stained with the Wigglesworth’s osmiumtetroxide-ethyl gallate method. While all neurons were traceable in each sample, resolution of this method was highly limited and only the gross neural backbones were reconstructed, producing schematic projection patterns. Our data is therefore the first account of a comprehensive, three-dimensional description of EPG/PEG cells in the central complex of all six studied species.

In all species, the overall morphology of EPG-like cells was near identical. The similar relative positions of the main neurite in the bundle connecting the protocerebral bridge and the fan-shaped body indicates a similar placement within the neural lineages that produce the columnar cells of the central complex and thus clearly suggests that these cells are homologous across the studied insects. EPG-like cell types therefore likely emerged over 400 million years ago.

Besides individual morphology and cell type identity we investigated the repertoire of EPG-like cells within each central-complex column as well as their overall projection patterns. Across holometabolous insects, we consistently found four copies of EPG-like cells per column, with the exception of the midline adjacent columns, which contained one copy less. The studied hemimetabolous species showed a near identical neural repertoire, only reduced by one EPG cell in each column. In locusts, one EPG-like cell type was present as a fourth neuron, but lacked an axon to the gall. Indeed, a set of four EPG-like cells (CL1a-d), with one cell lacking a gall projecting axon, was reported in locusts based on dye injected and Golgi stained examples (***Heinze and Homberg, 2008***). Four neurons per column were also found in the *Drosophila* connectome (***Hulse et al., 2021***). Overall, these highly consistent cell numbers indicate tight evolutionary constraints on evolvability of these neurons, in line with their essential role as head direction cells.

When analyzing the EPG neuron projection patterns, we found that these cells tile the ellipsoid body and each protocerebral bridge hemisphere into nine columns across all species. This pattern was first fully described in the fruit fly (***Hulse et al., 2021***), and recently confirmed in several species of bees and ants (***Sayre et al., 2021***; ***Sayre, 2022***). These sets of columns are connected in an identical way across all species, suggesting that the pattern of information flow between the ellipsoid body and the protocerebral bridge is highly conserved.

Interestingly, the original projection pattern analysis in locusts by ***Williams*** (***1975***) identified eight columns in the protocerebral bridge, missing the midline adjacent column, which is reduced in size in locusts. While an eight column arrangement was long considered the fundamental design of the central complex, our data show that the recent studies in flies and bumblebees indeed represent the default arrangement of this region. Intriguingly, in the fruit fly, the nine EPG-neuron defined columns are functionally arranged into the eight neural nodes of a ring attractor circuit, encoding the azimuthal space around the animal as a localized activity bump (***Seelig and Jayaraman, 2015***; ***Green et al., 2017***; ***Turner-Evans et al., 2017, 2020***; ***Hulse et al., 2021***).

While the full analysis of the head direction circuit was beyond the scope of this paper, we used automatic segmentation and synapse detection in the central complex of *Megalopta genalis* to identify the polarity of EPG and PEG neurons and to study their interconnectivity. We found that, in each column, there are three EPG neurons and one PEG cell, and that these neurons form a direct recurrent loop between the protocerebral bridge and the ellipsoid body. This configuration is identical to that of the fruit fly. Interestingly, the recurrent loop between EPG and PEG neurons is formed by direct, reciprocal connections in the bee, whereas the corresponding loop in the fly is closed indirectly via PEN neurons (***Turner-Evans et al., 2020***). While recurrent connections are a defining feature of the fly head direction circuit, our data provide first circuit-based evidence that different species might have evolved different solutions to solve the problem of computing a stable head direction signal. While highly conserved at the level of cell types, cell numbers and projection patterns, at the circuit level even this highly conserved network remains evolvable.

### Towards democratizing connectomics

In summary, we have developed and validated a novel workflow to support comparative connectomics projects at a scale where existing custom workflows, dedicated to specific full or near full-brain reconstruction projects, are impractical. By combining multi-resolution imaging and reconstruction, robust alignment, and reusable segmentation models, our approach aims at enabling small research groups to analyze neural circuits at the synaptic level. While large consortia remain essential for petascale datasets and full brain reconstruction at synaptic level, our method lowers the entry cost for hypothesis-driven studies of modular brain regions such as the one presented as a proof-of-concept.

Facilitating a community-driven connectomics approach, we have implemented the CAVE multiuser platform for proofreading. As this platform was originally designed for large full brain reconstruction efforts, running costs remain prohibitive when deployed for smaller datasets. To open this platform effectively for small projects, we have developed a communal cost sharing model, in which a single deployment of CAVE is shared by multiple research groups. This deployment is run by a commercial service provider (Tedore Interactive, Germany) and currently facilitates hosting datasets across five international research teams, substantially reducing project running costs for each group.

Overall, our entire workflow was designed to lower costs, reduce data volume, and develop standardized image alignment and segmentation routines that can be used off-the-shelf by any research group. We hope that this approach will enable research teams to conduct connectomics based neural circuit analysis, without the need to develop challenging customized workflows or to employ costly commercial solutions. By creating an accessible entry point to volume EM based circuit analysis, and thus broadening the field of connectomics, we hope to open the path towards answering numerous neurobiological questions that are currently out of reach.

## Methods and Materials

### Animals

Image data and neuron reconstruction presented in this paper were extracted from the CX of six insect species (Fig4a): the African praying mantis (*Sphodromantis lineola*), the Madeira cockroach (*Rhyparobia maderae*), the desert locust (*Schistocerca gregaria*), the European earwig (*Forficula auricularia*), a species of army ant (*Eciton hamatum*), and the tropical sweat bee (*Megalopta genalis*).

Praying mantis, cockroach, and desert locust specimen were collected from lab colonies and were processed directly after collection. Earwigs were caught in the wild near Lund (Sweden). Captured earwig specimens were kept in the fridge at 4° C with food ad libitum (leaves) until they were processed within 48 h. Sweat bees and army ants were collected in the tropical forest near the Smithsonian Tropical Research field station on Barro Colorado Island (Panama). *E. hamatum* workers were collected from foraging colonies and placed in ventilated vials containing cotton balls soaked with sucrose solution until being processed. To collect *M. genalis* sweat bees, a light trap was set up around twilight when these species are most active. Light traps consisted of a large white sheet illuminated by a light source containing UV light. Captured sweat bees were kept in a vial containing cotton balls soaked in sucrose and water solutions until processing within a week of capture.

### Histology for SBEM

For SBEM histology, brains were stained using a standard protocol as described below. We found great staining results using the method by ***Hua et al***. (***2015***), and therefore applied a slightly modified version of it to all insect species.

For the six insect species, began by anesthetizing the animal over ice for 3-5 minutes before removing the head. The brain was immediately dissected away from the head capsule in fresh fixative, removing the optic lobes, subesophageal ganglion, and the neural sheath whenever possible to aid with fixative permeation. Standard EM fixative contains 2.5% glutaraldehyde and 4% paraformaldehyde in 0.1M sodium cacodylate buffer. A fortunate dilution error made with *Megalopta genalis* led us to discover that using significantly lower concentrations yield far better tissue preservation. For all samples in this paper, we therefore used a fixative containing 0.75% glutaraldehyde and 0.75% paraformaldehyde in 0.1M sodium cacodylate buffer. After dissection, brains were fixed over night at 4° C. The following day, insect brains were rinsed 3 x 10 min in 0.1M sodium cacodylate buffer and then stored in the same buffer until further processing. Storage in buffer following fixation ranged between a few days and a few weeks, without any clear detriment to tissue quality.

For processing, brains were placed in a solution containing 2% osmium tetroxide in 0.1M sodium cacodylate buffer. The sweat bee, army ant, and cockroach were microwave treated (Biowave, Ted Pella) using two 1 min on/1 min off/1 min on steps at 150 W under vac, while the other brains instead underweat two rounds of 2min on/2 min off/2 min at 150 W under vac. Brains were then microwaved using 20 sec pulses at 250W three times, swirling the brains in solution between steps. The sweat bee, army ant, and cockroach were additionally left to rest for 20 sec between pulsations. Brains were next left to incubate in the same solution for 55 minutes at room temperature before being transferred to a new solution containing 1.5% potassium ferricyanide for the bee, ant, and cockroach, and 2.5% potassium ferricyanide for the praying mantis. The locust and earwig were instead transferred into a solution with 2.5% potassium ferrocyanide. Then, samples were microwave treated using the same pulse settings as described above. Bee, ant, and cockroach brains were left at room temperature for 1 hour.

All insect brains were next rinsed 4 x 5 min in ddH_2_O and then placed in a solution of 0.1% thiocarbohydrazide in ddH_2_O and left to incubate in the oven at 40° C for 45 min, omitting microwave pulsation steps. Following this, brains were transferred to fresh vials containing ddH_2_O and again rinsed 4 x 5 min with ddH_2_O. Tissue was then placed in 2% osmium tetroxide in ddH_2_O. Bee, ant, and cockroach brains were subjected to a single microwave pulsation step at 1 min on/1 min off/1 min (150 W) and left on the bench for 30 min. Other samples were instead subjected to twice microwave pulsation step at 2 min on/2 min off/2 min (150 W) and left on the bench for 30 min. Samples were next rinsed 4 x 5 min in ddH_2_O and transferred to 1% uranyl acetate in ddH_2_O. Samples were then left to incubate overnight at 4° C.

The next day, brains still in uranyl acetate solution were transferred into an oven and heated to 50° C where they were left to sit for 70 min. Samples were removed from the oven, cooled to room temperature, and then rinsed 4 x 5 min in ddH_2_O. A solution of lead aspartate was made by adding 0.066g lead nitrate to 10 ml of 0.03M aspartic acid in a falcon tube and pH adjusted to 5.5 with potassium hydroxide. The solution was heated at 60° C for an hour to aid dissolution. All insect brains were then soaked in lead aspartate solution for 45 min in the oven set to 50° C and then rinsed in ddH_2_O 4 x 5 min.

Lastly, brains were dehydrated through an ethanol series (20%, 50%, 70%, 90%, and 100%) at 15 min steps, transferred to acetone for 10 min, and then over the course of 2 hours embedded in Durcupan polymer (Sigma-Aldrich) at increasing ratios beginning from 3:1 (acetone:Durcupan), 1:1, 1:3, and finally twice into 100% Durcupan. Samples were then left to polymerize at 60° C for 48 hours.

### µCT-guided block trimming

#### Rough trimming and quality control

Once brains were stained and embedded, excess resin was trimmed into a cube using a razor blade under a stereo-microscope. The use of a saw or file is not advised as they may cause cracks in the resin due to vibrations. Blocks were shaped such that samples were roughly oriented with the brain standing vertically along its dorsal-ventral axis. From here on, we refer to the plane from which the SBEM scan will start as the imaging face and the opposite one as the stub face (which will be against the SBEM stub during image acquisition, Figure 1b-c). Cutting in the SBEM chamber will therefore go from the imaging face to the stub face. Each block was first mounted on a block against their imaging face using glue. The exposed stub face was polished with an ultramicrotome to produce a flat rectangular surface which facilitates the following fine trimming steps. The polished blocks obtained were scanned on their stub using a Yxlon FF35T micro-computed tomography (µCT) scanner as individual small resin pieces mounted with dental wax on a tall plastic stick. Neuropils of the insect brain are densely packed with stained neurites and are thus particularly salient under X-rays scans. This property allows for assessing structural condition and staining penetration, and identifying regions of interest.

To assess the quality of a sample, we first looked for cracks resulting from dissection or sample preparation. Insect brains contain networks of trachea that may be mistaken for damage but can be ignored when they look bilaterally symmetrical and similar across samples of the same species. Second, we identified the presence of density gradients along the sample which reflected poor staining penetration. Such issues typically appears as a light-to-dark gradient from the surface to the center of the brain. Samples were discarded when they presented cracks close to the region of interest or in the presence of density gradients which reflected poor penetration of staining agents. Blocks that passed quality control according to these criteria were selected for fine trimming.

#### Fine trimming with crosshair

µCT scans of the blocks passing quality control were opened in FIJI and imported in the plugin Crosshair (***Meechan et al., 2022***). Crosshair is an open-source FIJI plugin allowing to target a region of interest within resin-embedded samples by guiding ultramicrotome trimming of a block, based on a µCT scan. As described by ***Meechan et al***. (***2022***), the workflow consists in three major steps: the definition of a block and target plane, the alignment of the ultramicrotome knife, and the implementation of the ultramicrotome parameters computed by Crosshair. The block plane is one of the polished faces of the resin block, while the target plane defines the desired future block face and may be oriented differently to the current block face (fig1b). Both block and target planes can be defined by visual exploration and manipulation of the µCT scan using FIJI’s BigDataViewer (***Pietzsch et al., 2015***) and 3DViewer (***Schmid et al., 2010***). The alignment of the ultramicrotome knife ensures that the position of the knife matches that expected by Crosshair. Finally, Crosshair provides a set of ultramicrotome parameters such that the new block’s outer face becomes the target face after trimming. Crosshair was intended to be used with motorized ultramicrotome systems communicating with the plugin to automatically update the position of the ultramicrotome’s knife and sample arms. In absence of a motorized system, the solution provided for the RMC system was implemented with manual parameters input (see Materials and Methods section of ***Meechan et al., 2022***).

With the sample glued on its imaging side, the future stub face was defined as the target plane. Using FIJI’s 3DViewer, the sample was rotated until a suitable orientation for the region of interest was found. When possible, the sample was oriented such that the region of interest appeared roughly symmetrical and landmarks could be identified upstream of the region of interest along the axis of cut. Note that as much stained tissue as possible should be in contact with the metallic SBEM stub to maximize the dissipation of electrons from the sample. The stub face should therefore be placed past the end of the region of interest relative to the imaging face and should transect tissue. Ultramicrotome parameters were computed as per Crosshair’s instructions and the resin block was trimmed to the desired stub face using a diamond knife.

Once the stub face was trimmed to the desired orientation and depth, the block was carefully removed from its stub using a razor blade to slowly scrape off the glue holding it. The block was then glued in its final orientation against a Volumescope SBEM stub using conductive silver resin, and left to cure for 15 min. We ensured that silver resin did not cover the sides of the sample as it would shield it from X-rays. Another µCT scan of the mounted block was produced. This scan reflects the orientation of the sample as it will be in the chamber of the SBEM, and is used to trim the imaging face to a suitable depth and guide SBEM image acquisition.

The imaging face was identified by exploring the second µCT scan with FIJI. At this point, the imaging face does not require any transformation as it is parallel to the stub face and therefore the stub. To estimate the amount of resin to trim to reach the imaging face, the perpendicular distance from the imaging face to the block face was measured using FIJI. In cases where the block face was uneven, this was the greatest distance along the imaging face normal to its intersection with the block face.

To trim the block to the measured depth, the position of the knife was aligned to the base of the SBEM stub following the same workflow as proposed by Crosshair (***Meechan et al., 2022***). An empty resin block was mounted on an identical SBEM stub and the knife tilt was set to zero. The face of the empty block was polished, the knife was backed away, and the sample holder was rotated by 90°. The knife was then aligned to the sample’s polished face by only changing its tilt. With the knife such aligned, the empty resin block was replaced by our sample. The knife was carefully approached to the sample until it just began to cut, as indicated by the deposition of resin debris on the knife. Finally, the sample was trimmed to the depth measured at the previous step. This depth should account for a margin of error so as not to cut into the region of interest, as its accuracy depends on µCT resolution and trimming precision. Before placing the sample in the SBEM vacuum chamber, silver resin was applied to the sides of the block and left to cure to increase sample conductivity.

### Image acquisition

Volumetric image stacks were acquired with a VolumeScope SBEM using a VolumeScope directional back-scatter detector (VS-DBS; Thermo Fisher Scientific) and MAPS 3 software (Thermo Fisher Scientific). Samples were cut at 50 nm thickness and imaged using a landing beam energy of 2 kV, beam current of 0.1-0.2 nA, and pixel dwell time of 1-3 µs. Beam current and pixel dwell time were chosen for each scan based on their tendency to charge. High beam current and short pixel dwell time were favored when possible. We imaged under low vacuum conditions (10 Pa) to limit charging. Where multiple tiles had to be used for large fields of view, we defined an overlap of 15% of tile shape.

We used a multi-resolution approach consisting of a large field of view capturing the whole CX at cellular resolution (30-40 nm) and smaller overlapping fields of view capturing compartments of the CX at synaptic resolution (8-12 nm). Pixel size in XY was determined to account for different brain sizes between animals. The cellular resolution images must allow the reconstruction of neuron backbones and large branches, while small neurites and synaptic sites must be resolved in synaptic resolution images. We found that a resolution of 8-12 nm typically allows the identification of pre-synaptic densities but not gap junctions. Synaptic resolution compartments were carefully placed to follow the projection patterns of a few of the repeating computational units of the CX. Throughout the acquisition of a sample, we used the µCT scan of the mounted sample acquired in the previous preparation step to guide the placement of imaging tiles.

### Image processing

SBEM produces thousands of 8 bit greyscale images in TIF format, tiling the region of interest. These must be aligned in three dimensions to produce contiguous images stacks allowing the reconstruction of neurons across all regions acquired. We wrote a custom python pipeline for image alignment, relying on Google Brain’s SOFIMA package (***Januszewski et al., 2024***) which uses GPU-accelerated optic flow estimation and elastic mesh optimization to non-linearly align EM images. We start by fully aligning the cellular resolution image stack, which will be used as a frame of reference to all synaptic resolution stacks in the same dataset.

#### Input data

The MAPS 3 software produces one directory per tileset, a group of image tiles organized on a rectangular grid with overlap between neighbors. All images in a directory also share the same resolution, imaging parameters, and shape. Their position in the grid and slice number are contained in their name, and a text file per slice contains their resolution. This information is recovered automatically, and tiles belonging to the same tileset are organized into tile maps, where images require only rigid transformations to be coarsely aligned to each other. This notably facilitates XY alignment using SOFIMA, which provides a function to align tile maps requiring tiles to have the same shape at the time of writing (version: 20240120.dev36+g5c5642c). The code responsible for organizing images was written to match the file structure produced by ThermoFischer’s MAPS 3 software’s output. We provide a guide to modify the functions identifying directory structure such that it can be adapted to other microscopes.

#### Alignment planning steps

At the start of a new project, one sample image is selected from each of the directories that were detected by the previous step and loaded into a local neuroglancer viewer. The user is then prompted to examine sample images, to determine whether a directory’s tile should have their histogram inverted such that it matches standard EM conventions (i.e. electron-dense regions should appear dark). One configuration file is consequently written per directory, and contains information about image tile paths and placement, and instructions for histogram inversion determined from user input. One main configuration is additionally written which contains paths to all configuration files for a project, as well as information that apply to all directories such as output paths and alignment parameters. Alignment parameters are either determined from default values or user input. Images can optionally be pre-processed before XY alignment using a gaussian filter for denoising and CLAHE for contrast adjustment as per OpenCV implementations (***Bradski, 2000***).

For our datasets, images were consistently denoised using a gaussian filter with sigma 1 and kernel size 3 x 3, and contrast-enhanced using CLAHE with clip limit 2 and tile grid size 10 x 10.

A similar planning step occurs between XY and Z alignment. Aligned tile maps are first examined for overlap if multiple images exist for the same slice index. Overlap is determined using SIFT keypoints detection, and is deemed valid if enough keypoint matches of good quality are found between images. In such case, a configuration file is written to inform the fusion of overlapping aligned tile maps. Before Z alignment, aligned tile maps are compared to find the project’s root slice, to which all subsequent slices will be aligned. The root slice is the first one containing a single image, as opposed to slices where multiple tile maps exist without overlap. One configuration file per tile map is written, containing parameters for Z alignment and order of alignment starting at the root slice. Configuration files will be used for Z alignment and can be modified on an individual basis to provide finer control over alignment of different intervals of image data along the Z axis. Slice indices can optionally be provided to indicate slices of poor quality that should be ignored for alignment purposes. Images are still warped using transformations computed for their neighbor’s, but they do not contribute to other images’ alignment computation.

#### Alignment with SIFT and SOFIMA

Throughout this pipeline, we define a coarse alignment consisting in rigid or affine transformations, and a fine alignment consisting in non-linear elastic transformations. Although linear transformations can provide satisfactory alignment to some extent, elastic alignment is necessary to correct for non-linear deformations caused by the back-scatter detector, most notably occurring at the edges of the field of view. Such artifact can make neuron reconstruction difficult within regions of overlap between images. For coarse alignment, we use SOFIMA (***Januszewski et al., 2024***) to compute rigid transformations for XY alignment of a tile map, and we use the OpenCV (***Bradski, 2000***) implementation of SIFT keypoints detection to compute affine transformations outside of tile maps. Fine alignment is systematically done using SOFIMA to compute optic flow within overlapping patches of data at regular intervals. Optic flow is then regularized with an elastic mesh to create smooth and minimal deformation.

#### XY-alignment

We start by aligning the cellular resolution image stack in XY. Images are loaded and organized in tile maps based on configuration files produced during the planning phase. Tile maps are first coarsely aligned with rigid transformations before fine elastic alignment, both using SOFIMA, producing one image stack per tile map. If tile maps are found to overlap after XY alignment is completed, they are coarsely aligned to each other with OpenCV’s SIFT, and finely aligned with SOFIMA.

#### Z-alignment of cellular resolution images

Once all overlapping tile maps are stitched, images are aligned along the Z axis to provide smooth transition between slices. Alignment starts with a root slice which will serve as a reference for the whole dataset. Each subsequent slice is then aligned to its last processed neighbor. That is, if the root slice is found at an index higher than 0 along the Z axis, slices at lower indices will be aligned to their direct neighbor with a higher index, while slices with indices higher than the root will be aligned to their neighbor with a lower index. For each slice, images are first coarsely aligned with SIFT before using SOFIMA for fine alignment. Missing slices are ignored and will result in a black slice in the final stack. They do not contribute to the final alignment.

#### Synaptic resolution image alignment

For synaptic resolution images, we start with XY alignment as described above. Once all tile maps have been aligned within the XY plane, they are aligned to the cellular resolution image stack acting as a frame of reference on the same slice. If a cellular resolution slice is missing, or if sampling along the Z axis was coarser for the cellular resolution stack, the closest slice is chosen as a reference. As the cellular resolution stack was already aligned along the Z axis, it is unnecessary to align neighboring slices within the synaptic resolution stacks.

#### Output data

Configuration files are saved as JSON files at each step to ensure reproducibility. Wherever iterating through images is necessary, progress is logged to a MongoDB database to ensure that it can be resumed in case of interruption. Progress can also be wiped for arbitrary intervals and steps, such as for re-computing alignment with new parameters for a given region. Alignment parameters and metrics are also logged to facilitate troubleshooting. All arrays are written in the Zarr format, which is particularly suited for parallel task computing and will be used in all downstream steps of the image processing pipeline (such as neuron segmentation). Aligned images, affine transformation for coarse alignment, and optic flow estimations and relaxed mesh for fine alignment are written to file after XY and Z alignment. A copy of the output of Z alignment is written in a downsampled form to facilitate the examination of aligned image data. Optic flow estimations are saved so that mesh relaxation may be repeated if final alignment is not satisfactory. This may notably happen when image data varies in quality over long image acquisition bouts, if charging occurs, or if slices get missing. Configuration files, affine transformations, and relaxed mesh are light-weight data that can together be used to deterministically reproduce an aligned image stack from raw data, while reducing computation times to warping steps only.

### Estimation of time gained

Estimation of time gained by our imaging strategy was computed using python. For each dataset considered, all raw data images were listed with their shape, resolution, and the dwell time used to acquire them.

Consider a tile map made of a grid of *g*_*x*_ by *g*_*y*_ images and *n*_*z*_ slices. Each image has the same *n*_*x*_ by *n*_*y*_ pixels shape for a given tile map. All images belonging to the same imaging run were acquired using the same dwell time *t*_dwell_. Therefore, we can calculated acquisition time for a tile map as:

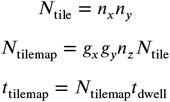

where *N*_tile_ is the number of pixels for a tile, *N*_tilemap_ the total number of pixels in a tile map, and *t*_tilemap_ its total acquisition time. Effective total imaging time *t*_total_ is therefore estimated as:

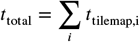

We then calculated a hypothetical acquisition time in a scenario where all image data is acquired at synaptic resolution. Only cellular resolution tile maps should be considered as their area covers the entire region of interest. The area *A*_tilemap_ of tissue acquired corresponds to the area of tissue covered by tile maps, taking into account the overlap between neighboring tiles:

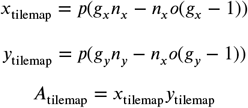

where *x*_tilemap_ and *y*_tilemap_ are the width and height of tissue covered by a tile map in world units, *p* the pixel size used to acquire that tile map, and *o* the overlap between tiles is expressed as a percentage of their shape as is defined in the MAPS 3 software used for image acquisition. We used an overlap of 15% of tile shape for our calculations. In an hypothetical scenario where the whole region of interest was acquired at synaptic resolution with a pixel area *P*_highres_, acquisition time *t*_highres_ would be estimated as:

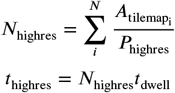

For each dataset, the pixel size in the hypothetical acquisition scenario was the same as the one used for synaptic resolution compartments of that dataset (or the most used one when resolution varied). Cutting time would not vary from the real acquisition and could therefore be ignored. Note that these calculations underestimate the number of pixels resulting from the hypothetical acquisition scenario, as we ignore overlap between tiles. Using higher resolution with the same tile shape would increase the number of tiles and therefore the total area of overlap, resulting in a larger total number of pixels *N*_highres_ and thus a longer acquisition time that the one estimated here. Tiles could in theory be made larger but this would come at the cost of distortions at the periphery of the field of view.

### Neuropil manual segmentation

Neuropils were segmented using Amira segmentation editor or the open-source software 3Dslicer (https://www.slicer.org/; ***Fedorov et al., 2012***). The neuropils for *Megalopta genalis* and *Eciton hamatum* were segmented by downsampling the cellular resolution image data to 0.4 x 0.4 µm pixel scale in XY using FIJI. Neuropils were manually segmented by painting within the limit of the neuropil at regular intervals in the xy, xz, and yz planes. The Wrap Tool was then used to interpolate planes and compute a smooth surface around the neuropil. Neuropils for other species were segmented using 3Dslicer. Image data was first downsampled to 0.4 x 0.4 µm pixel scale in XY using OpenCV, and written to an 8 bit 3D TIFF file which could be opened by 3Dslicer. As with Amira, neuropils were manually segmented by manually painting at regular intervals. The resulting sparse segmentation was interpolated to compute a smooth volume of the neuropil.

### CATMAID implementation and manual tracing

CATMAID was deployed as per instructions that can be found at https://catmaid.readthedocs.io/en/stable/. CATMAID is hosted on three private servers: the app, database, and image data are hosted separately. Synaptic resolution image data was notably served to CAVE directly to avoid unnecessary duplication. Backend deployment and management were handled by a commercial IT support (Tedore Interactive). Skeletons were traced manually using CATMAID (***Saalfeld et al., 2009***), a collaborative and open-source web-based annotation tool. Skeletons are traced by scrolling through image stacks and placing nodes within neuron boundaries to follow the path of its branches. A skeleton provides a minimal representation of a neuron’s arborization and cable length, but does not capture precise morphology. We manually traced the backbone of all neurons of the CX between and within neuropils approximately until reaching the second branching point away from the backbone, or until neurites could not be resolved confidently within cellular resolution image stacks. Backbones therefore bridge the gap between synaptic resolution image stacks which were segmented automatically.

### Automatic segmentation

#### Model architecture

Neurons within synaptic resolution image stacks were segmented using a 3D U-net artificial neural network with an architecture described as Multi-Task Local Shape Descriptors (MTLSD) in ***Sheridan et al***. (***2023***). This model is designed to predict nearest-neighbor affinity (NNA) and local shape descriptors (LSDs). NNA provides a probability for neighboring voxels to belong to the same label class, while LSDs provide local shape metrics and are used as an auxiliary task to improve the model prediction of NNA.

#### Segmentation pipeline

We produced segmentation using the lsd python package (***Sheridan et al., 2023***). Synaptic resolution image data was passed through the network to compute NNA, while LSDs were not saved as they are not used for downstream steps. The resulting NNA prediction array was post-processed following the method described in ***Sheridan et al***. (***2023***). Supervoxels are generated using a seeded watershed algorithm while storing their center of mass as a node in a region adjacency graph (RAG), with an edge between every pair of touching node. Edges are agglomerated by hierarchically merging them in order of decreasing affinity based on the predicted NNA, to compute a merge score from 0 to 1 weighing the edges of the RAG. Final segmentation can be obtained from the resulting segmentation graph by discarding all edges above a given merge score threshold and relabeling supervoxels according to the connected component they belong to.

#### Training and evaluation

Five 5x5x5 µm (100 x 500 x 500 pixels) volumes were selected as ground-truth out of the *Megalopta genalis* synaptic resolution dataset. Neurons and mitochondria within these volumes were manually segmented using Napari (***Sofroniew et al., 2025***) by an expert annotator. Neurons and mitochondria were segmented exhaustively such that they were all labeled within the ground-truth volumes. Where the identification of tissue boundaries was ambiguous (for thin branches or glia), neurons were segmented with a conservative approach favoring splits over merges. Both neurons and mitochondria were segmented in the ground truth, based on the intuition that mitochondria can provide additional contextual information to the model as they cannot cross neuron membranes. Models were trained using gunpowder (https://github.com/funkelab/gunpowder/tree/main) and PyTorch (***Ansel et al., 2024***) for up to one million iterations per training run, while saving a checkpoint model state every 50000 iterations. At each iteration, 36 x 124 x 124 pixel volumes of raw and ground-truth segmentation were selected randomly from ground-truth volumes and were augmented with gunpowder. Augmentation parameters were optimized with a random search setting parameter values before each training run.

Models were evaluated by computing variation of information (VOI) between segmentation and ground-truth skeletons, as described in ***Sheridan et al***. (***2023***). Two 10 x 10 x 10µm test volumes were selected from the *Megalopta genalis* synaptic resolution image data. These volumes were chosen to be visually representative of the majority of the image data for this dataset. Within test volumes, skeletons were manually traced using CATMAID and reviewed by expert annotators. Each checkpoint of each model was used to produce segmentation of the testing data. The segmentation graphs such obtained were iteratively thresholded with values from 0 to 1 with steps of 0.02, and VOI was computed by comparing the resulting segmentations to test skeletons. The best model was the one obtaining the lowest VOI score, signifying low discrepancy between skeletons and segmentation. We used the best model obtained from a batch of 20 training runs for all segmentation presented in this paper. Merge score threshold was chosen for each synaptic resolution segmentation volume by visual examination to favor false splits over false merges.

### Segmentation proofreading

Segmented data contains false merges (fragments from different neurons incorrectly joined) and false splits (fragments of the same neuron incorrectly separated). To correct these errors we deployed CAVE (***Dorkenwald et al., 2025***), a collaborative proofreading and annotation web-based platform for neuron segmentation.

As CAVE requires segmentation data to be ingested in a chunkedgraph (***Dorkenwald et al., 2022***) (CG) representation, we wrote custom python scripts to convert our segmentation produced as per ***Sheridan et al***. (***2023***) into the CG-compatible graphene format: each supervoxel is assigned a unique 64-bit label encoding its CG layer ID, chunk coordinates, and a segment ID unique within its chunk.

Parallel computations are performed chunk-wise with daisy, and a MongoDB is used to store intermediate mapping data. First, a supervoxel array is translated chunk by chunk into the graphene format and subsequently uploaded to a Google bucket in the cloudvolume format. For each chunk, we store a dictionary mapping original supervoxel IDs to CG IDs for the later translation of edges. In a second step, edges are retrieved from MongoDB for each chunk and translated into CG IDs using the corresponding dictionary mapping. Edges are classified into “in-chunk”, “between-chunk”, and “cross-chunk”. In-chunk edges connect supervoxels within the same chunk (with the same chunk ID). Between-chunk edges connect supervoxels across chunk boundaries (with different chunk IDs). Cross-chunk edges connect two CG supervoxels resulting from splitting one original supervoxel across chunk boundaries. CG edges are weighted by an affinity score between 0 and 1, with high values indicating high affinity, thus corresponding to the contrary of the merge score used throughout the segmentation pipeline (1 - merge score). Edges sorted in in-chunk, between-chunk, and cross-chunk are uploaded along with the corresponding affinity score to a Google cloud storage bucket in protobuf format.

Finally, the initial agglomeration state of the CG is computed. For each chunk, we construct a graph of CG edges and those of its direct neighbors. This graph is thresholded by discarding edges whose affinity score is above a user-defined threshold value, before computing connected components. We choose a threshold that produces a satisfying level of agglomeration while limiting false merges which are more labor-intensive to correct than false splits. The resulting agglomeration will be the initial state of segmentation for the uploaded volume to be proofread.

In our image data, image data near the boundaries of neuropils often contain large neuron branches with burst membranes, resulting in many false merge errors which can spread dramatically. To mitigate the effect of these artifacts on the initial state of agglomeration, we choose to split connected components at chunk boundaries. This results in segmentation fragmented at regular intervals, mostly requiring merge operations but limiting the number of large merge artifacts which are difficult and time-consuming to split.

### Proofreading strategy

We reconstructed neurons at two levels of resolution. We started by tracing the backbones of all neurons of interest within the cellular resolution image stack hosted in CATMAID. This provides a minimal representation of projection patterns which is generally sufficient to identify cell classes in the insect brain. Backbones also bridge the gap between synaptic resolution compartments that may not be overlapping with each other.

To reconstruct neurons in synaptic resolution volumes, we combine CATMAID skeletons and CAVE segmentation. We developed a web application which provides a means for users to query neurons from CATMAID into the CAVE ecosystem to guide segmentation proofreading. Users may search neurons in the linked CATMAID database using their name or annotations. One or more selected skeletons are matched to CAVE segmentation by listing segments IDs found at skeleton nodes coordinates. If a match is found, skeletons are converted to the neuroglancer format and displayed alongside corresponding segment IDs in the neuroglancer viewer used by CAVE.

Proofreaders therefore start with a list of automatically queried segment IDs corresponding to neurons of known identity, which they can expand by finding missing branches. Proofreaders begin by splitting segments when necessary before merging all the remaining clean fragments into one contiguous object representing the neuron.

### Synapse detection

Automated prediction of synapses was performed using the synful python package (***Buhmann et al., 2021***). Model architecture was the same as described by ***Buhmann et al***. (***2021***) and is designed to predict pairs of pre- and post-synaptic sites (Fig.4). Nine 5x5x5 µm (100x500x500 pixels) cubes extracted from the synaptic resolution image stacks and used to train 16 models. Training was done using gunpowder (https://github.com/funkelab/gunpowder/tree/main) with data augmentation. Models were evaluated following the method used by (***Buhmann et al., 2021***). Synapses were assigned to skeletons traced by an expert annotator within a testing volume of the same size as the ground-truth but not overlapping with it. Models were evaluated based on recall compared to an expert-annotated cube not included in ground truth. Scoring was optimized to reduce false positives while ensuring high recall. The best performing model was selected and used to compute synapses shown in figure 4. On representative samples from all three synaptic resolution volumes acquired for *Megalopta genalis* in the PB, FB, and NO, this model performed with an F-score of 0.662, 0.749, and 0.636 respectively. All F-scores are comparable to those achieved by ***Buhmann et al***. (***2021***) with image data of the fruit fly’s brain. A threshold was applied to all synapse predictions to remove low-confidence predictions.

### Data analysis

The phylogenetic tree in figure 4b was created based on data from ***Rainford et al***. (***2014***) and using the platform Interactive Tree of Life (iTOL; ***Letunic and Bork, 2024***). All visuals showing neuron morphology in figure 4a were created using the software Blender (https://www.blender.org/). Circuit graphs in figure 4d-e were made using the networkx python library using automatically detected synapses. Any connection with three or less synapses were discarded for connectivity graphs. Any autapse (synapse connecting the same neuron to itself) were considered false positives and were discarded.

## Code availability

The code for image alignment is available at https://github.com/Heinze-lab/EMalign and was adapted from Google research’s SOFIMA (original code at: https://github.com/google-research/sofima, ***Januszewski et al., 2024***). Code and models for 3D U-Net training are available https://github.com/Heinze-lab/EMtrain. The code for the segmentation pipeline is available at https://github.com/Heinze-lab/EMsegment. Both training and segmentation code are are adapted from local-shape descriptors (https://github.com/funkelab/lsd, ***Sheridan et al., 2023***). The code for synapse detection is available at https://github.com/Heinze-lab/synful_312 and based on work by ***Buhmann et al***. (***2021***) (https://github.com/funkelab/synful). The code for uploading segmentation to a CAVE instance is available at https://github.com/Heinze-lab/SegToPCG. The code for the catmaid-CAVE app is available at https://github.com/ktedore/catmaid-neuroglancer-api (backend) and https://github.com/ktedore/catmaid-neuroglancer (frontend).

## Acknowledgments

We thank the facilities, and the scientific and technical assistance, of the Microscopy Australia Facility at the Centre for Microscopy and Microanalysis (CMM), The University of Queensland. We are especially grateful to Roger Wepf, Kathryn Green, Robyn Chapman, and Rick Webb for valuable insights and access to the EM facility. We thank Robert Parton for collecting and providing live bull ants. We are grateful to Vivek Nityananda for providing the lab specimen of praying mantis and the facilities to perform dissections. Madeira cockroach and desert locust were provided by the Uwe Homberg lab. We thank Vanessa Althaus for performing the dissection of the cockroach and locust brains, and Martina Fromandi for arranging sample shipment. We thank Niklas Wahlberg for capturing and providing wild earwigs. We are grateful to the CAVE team for continuous support with implementing and troubleshooting our CAVE deployment. M.E.S. was funded in part by an International Cotutelle Macquarie University Research Excellence Scholarship (iMQRES 2019060), and by Australian Research Council grant DP220102836. S.H was supported by the European Research Council (ERC), under the Horizon 2020 and Horizon Europe frameworks (grant agreements 714599 – BrainInBrain and 01044220 – EvolvingCircuits), the European Innovation Council (EIC, grant agreement 101046790 — InsectNeuroNano), the Swedish Research Council (Vetenskapsrådet, grant agreement 2018–04851) and the Crafoord Foundation (grant agreement 20200709). Views and opinions expressed are however those of the author(s) only and do not necessarily reflect those of the European Union or the European Innovation Council. Neither the European Union nor the European Innovation Council can be held responsible for them. The funders had no role in study design, data collection and analysis, decision to publish, or preparation of the manuscript.

## Authors contributions

**Valentin Gillet**: Conceptualization, Methodology, Software, Investigation, Data curation, Writing — Original Draft, Writing — Review and Editing, Visualization; **Marcel E. Sayre**: Conceptualization, Methodology, Investigation, Data curation, Writing — Review and Editing; **Griffin Badalamente**: Methodology, Software, Investigation; **Nicole L. Schieber**: Methodology **Kevin Tedore**: Software; **Jan Funke**: Supervision; **Stanley Heinze**: Conceptualization, Methodology, Resources, Data curation, Validation, Writing — Review and Editing, Supervision, Project administration, Funding acquisition

---

**Figure 1—figure supplement 1.**
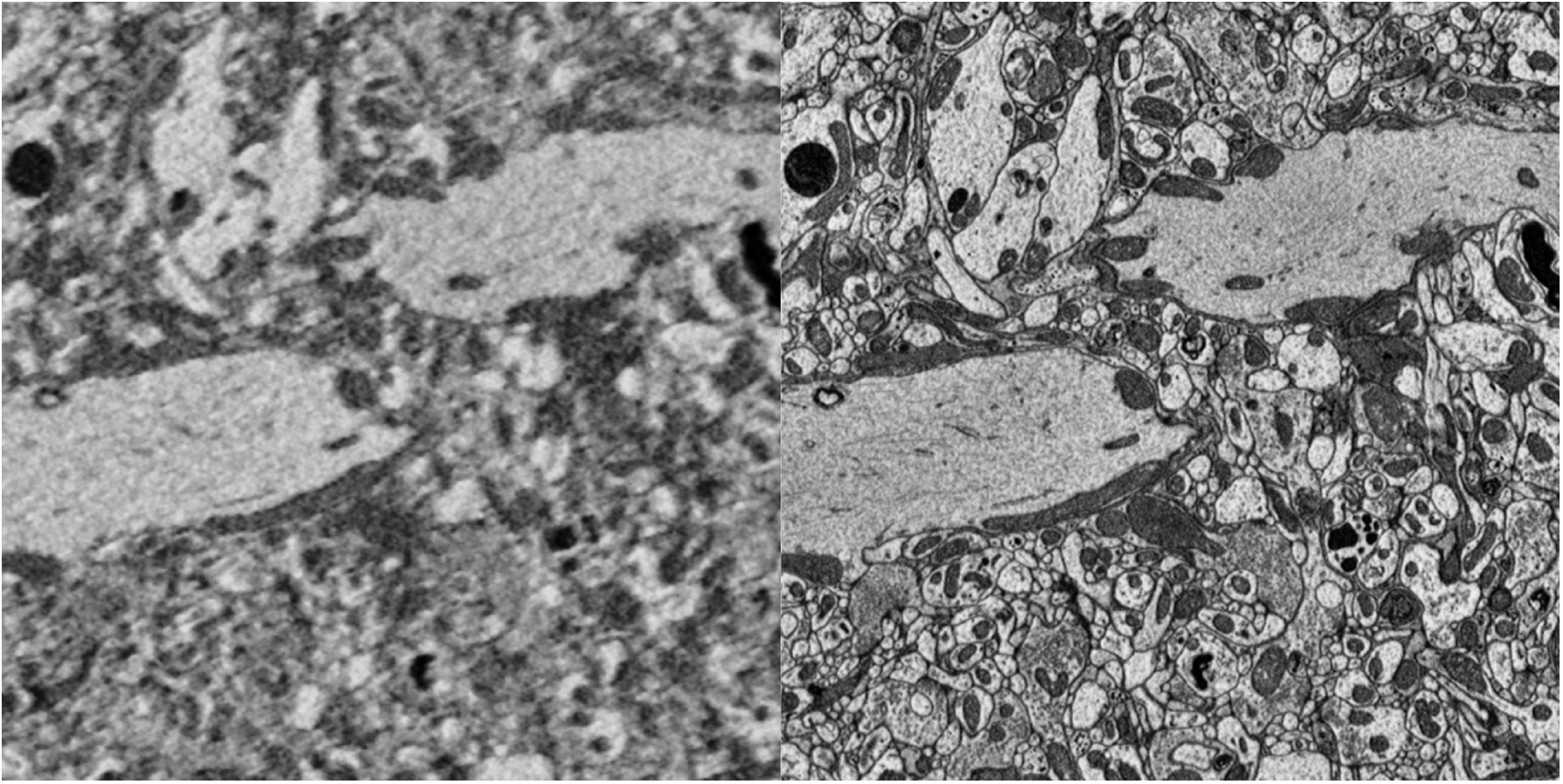
Zoomed in comparison between cellular and synaptic resolution image data. Images extracted from the same region of the protocerebral bridge of *Megalopta genalis*, illustrating the correspondence between cellular resolution (40 x 40nm pixel size, left) and synaptic resolution (10 x 10nm pixel size, right)

**Figure 2—figure supplement 1.**
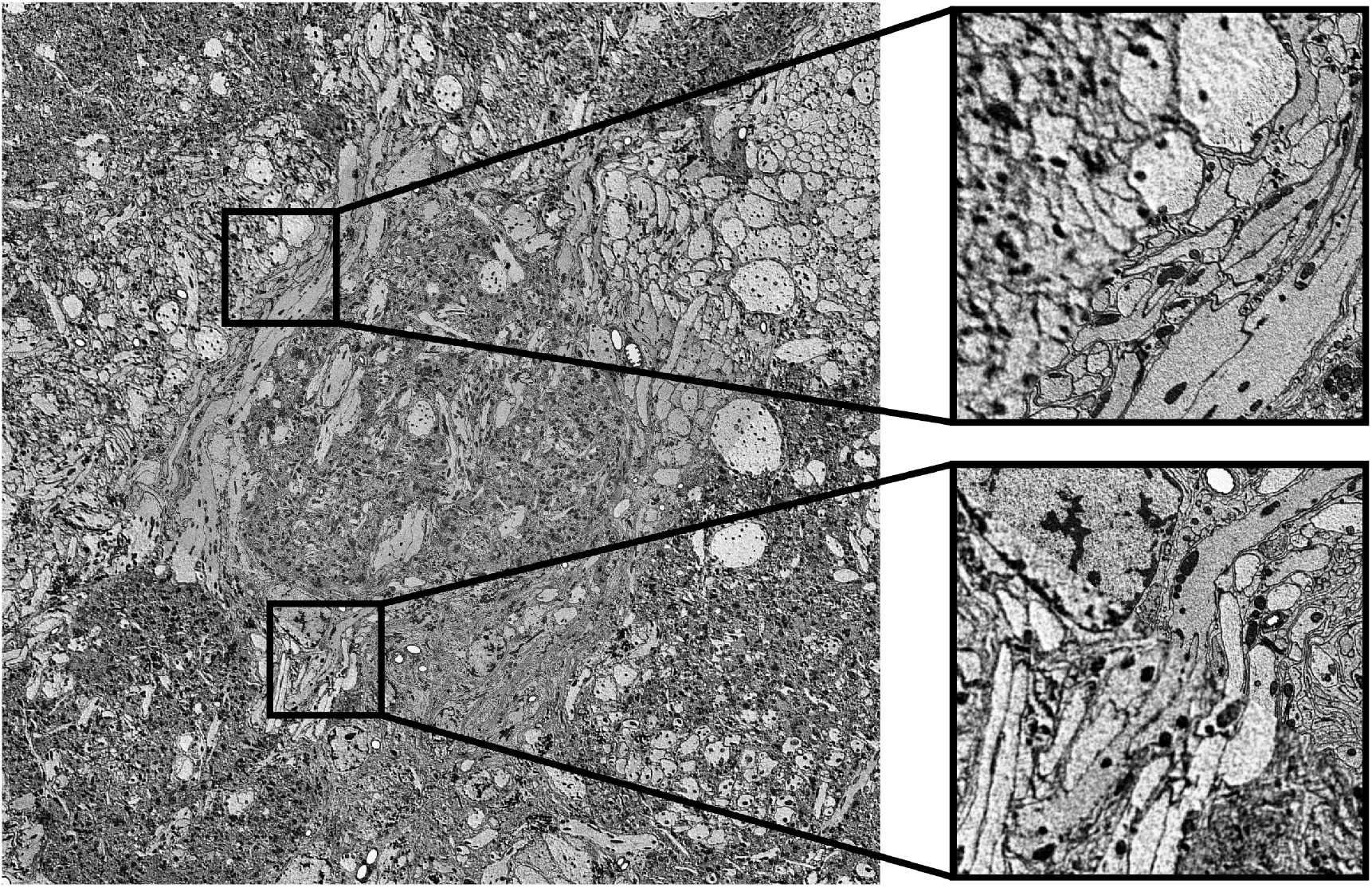
Alignment of synaptic resolution data to cellular resolution images.

**Figure 2—figure supplement 2.**
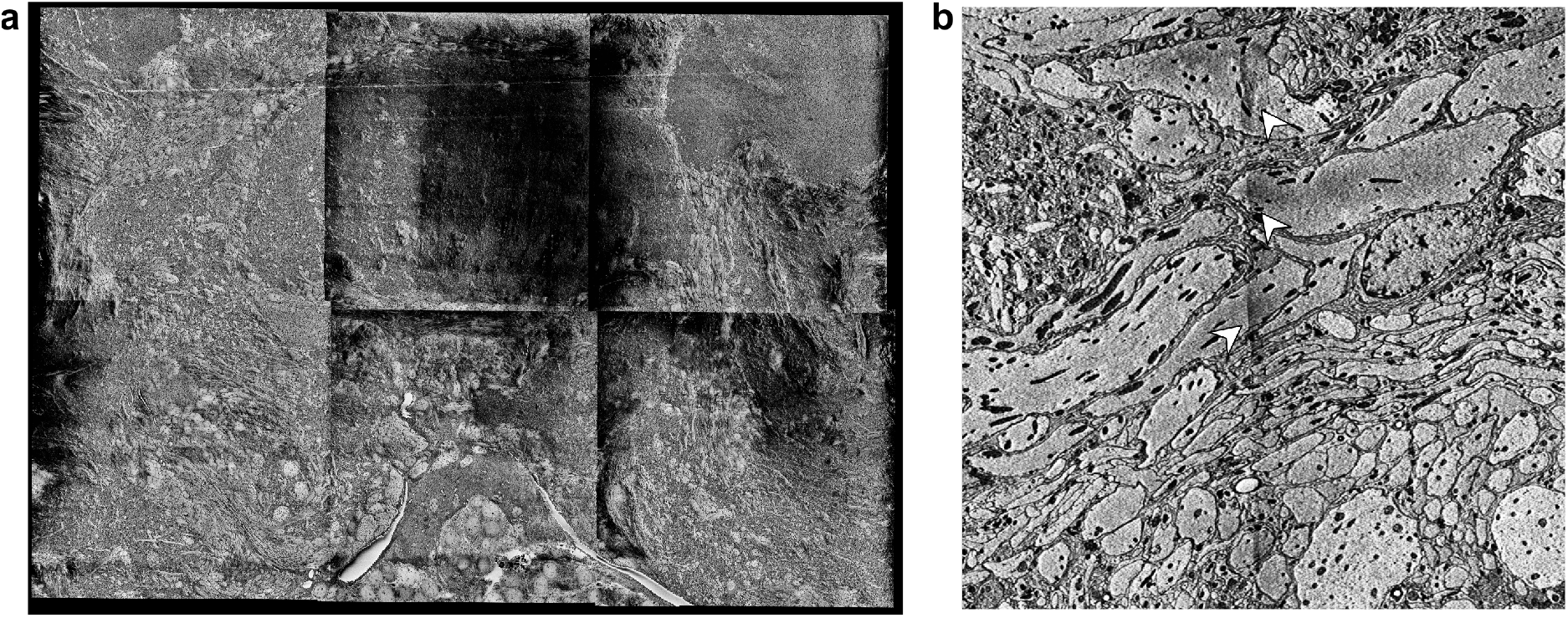
Example images showing charging artifacts. Images of the locust dataset illustrating the effect of charging on the image data. Panel a shows dramatic charging that made alignment impossible. Alignment was possible in panel b despite contrast difference because charging was minimal.

**Figure 3—figure supplement 1.**
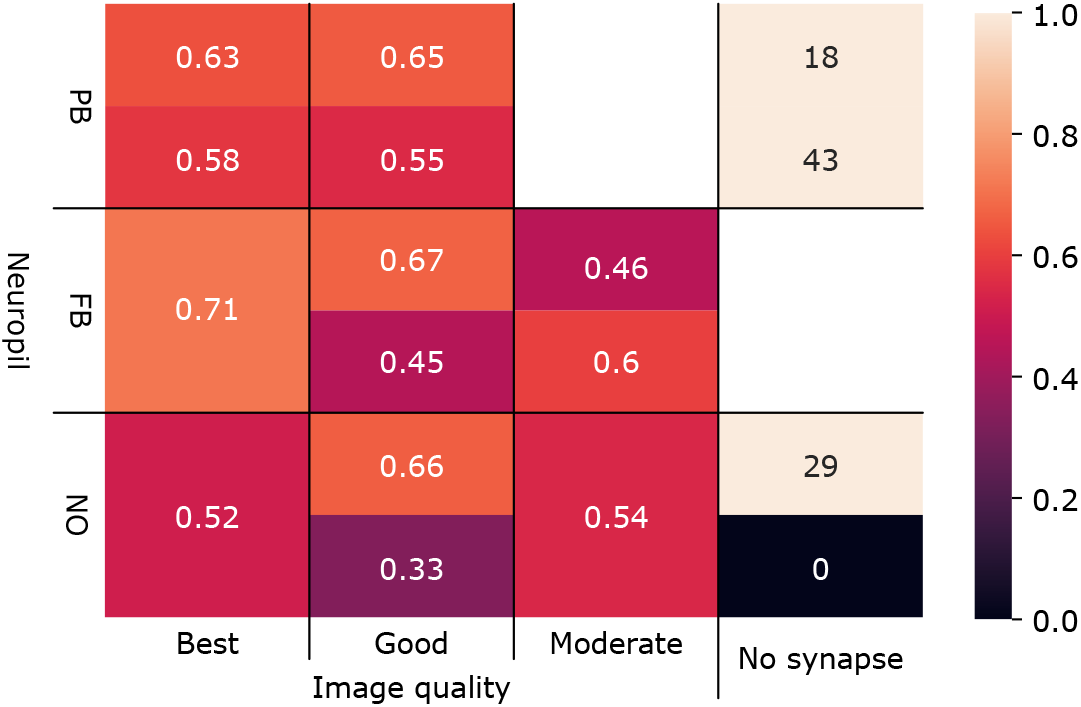
F-scores for synapse prediction. F-scores were computed for each neuropil dataset in *Megalopta genalis* for three kinds of image quality decreasing from “Best” to “Moderate”. Some images with no synapses were presented and expected to not have any prediction.

**Figure 3—figure supplement 2.**
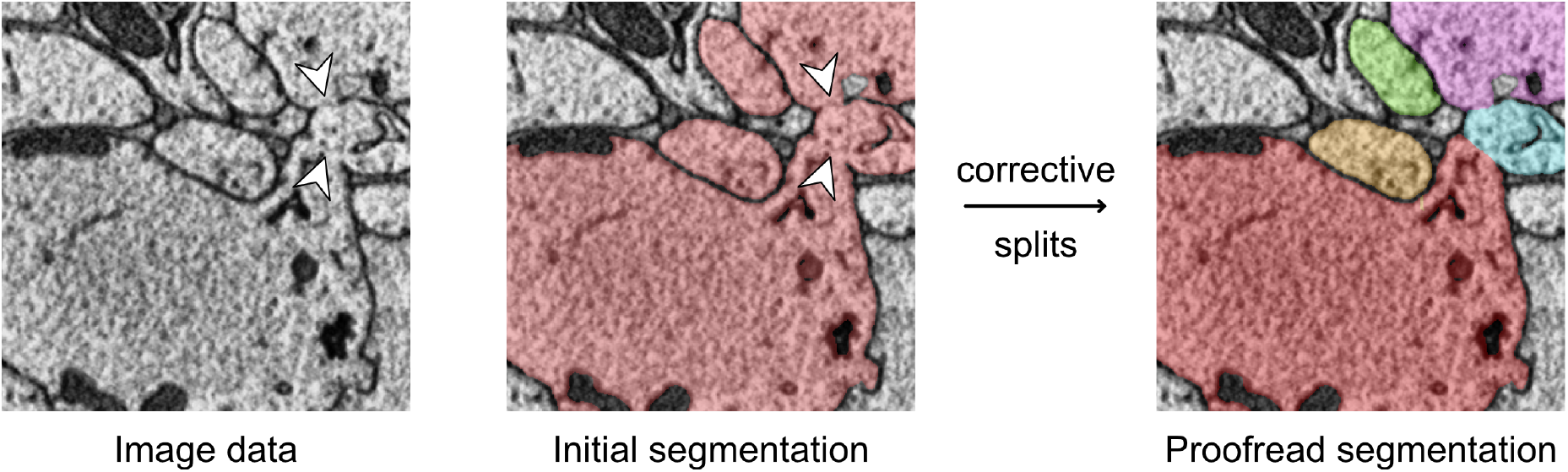
Example of a hole in cell membrane causing false merges in image segmentation. Image showing holes in the membranes of three neurons (arrows) that caused a false merge with automatic segmentation. Multiple neurons are initially assigned the same label as shown by their color. Some neurons do not appear to have holes but are likely connected outside of the region shown. After performing corrective splits via CAVE, the segmentation accurately reflects neuron morphology.

**Figure 4—figure supplement 1.**
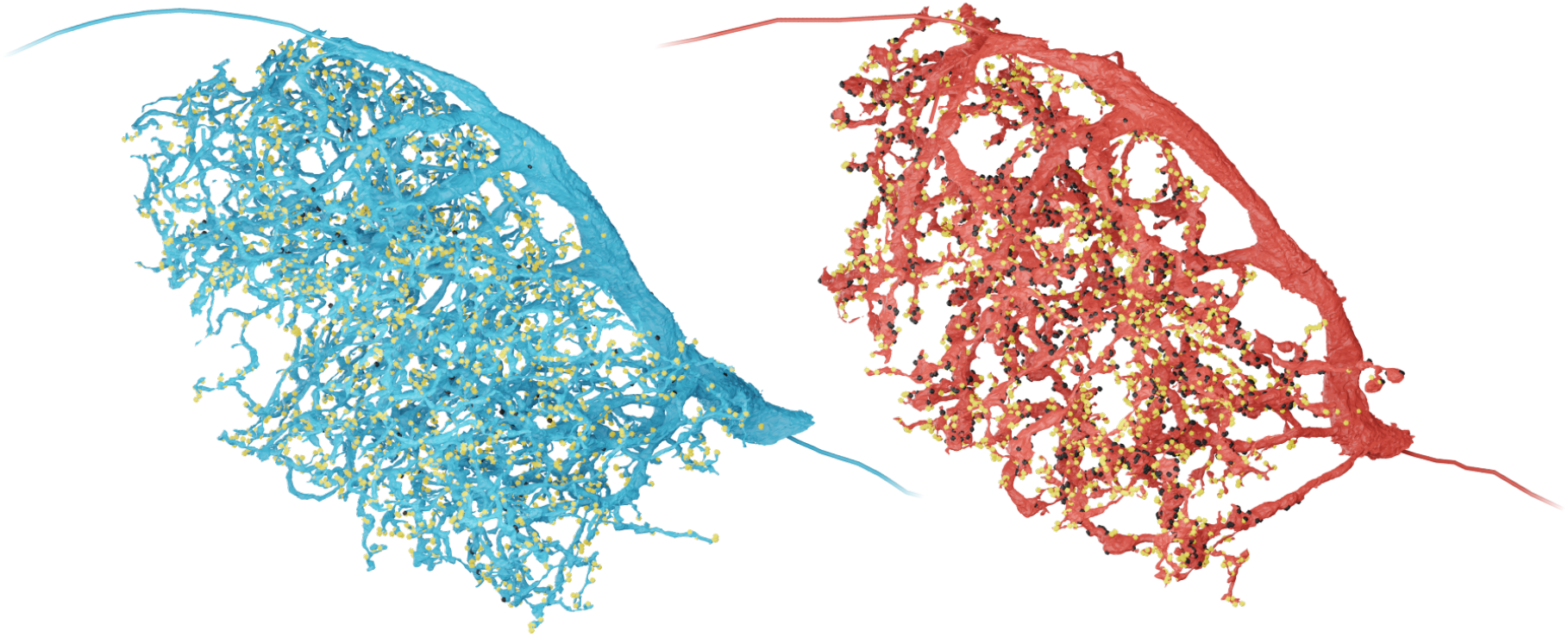
Side-by-side comparison of EPG and PEG neuron in the protocerebral bridge. Reconstruction of protocerebral bridge branches of an EPG (right, red) and a PEG (left, blue) neuron. Pre-synaptic sites are shown in black, post-synaptic sites in yellow.

